# Vexitoxins: a novel class of conotoxin-like venom peptides from predatory gastropods of the genus *Vexillum*

**DOI:** 10.1101/2022.01.15.476460

**Authors:** Ksenia G. Kuznetsova, Sofia S. Zvonareva, Rustam Ziganshin, Elena S. Mekhova, Polina Dgebuadze, Dinh T.H. Yen, Thanh H.T. Nguyen, Sergei A. Moshkovskii, Alexander E. Fedosov

**Affiliations:** Federal Research and Clinical Center of Physical-Chemical Medicine, 1a, Malaya Pirogovskaya, Moscow, 119435, Russia; A.N. Severtsov Institute of Ecology and Evolution, Rus. Acad. Sci. Leninsky prospect, 33, Moscow, 119071, Russia; Institute of Bioorganic Chemistry, Rus. Acad. Sci. Miklukho-Maklaya st, 16/10, Moscow, 117997, Russia; Russian-Vietnamese Tropical Research and Technology Center, Coastal Branch, 30 Nguyễn Thiện Thuật, Nha Trang, Vietnam; Pirogov Russian National Research Medical University, 1, Ostrovityanova, Moscow, 117997, Russia

## Abstract

Venoms of predatory marine cone snails (the family Conidae, order Neogastropoda) are intensely studied because of the broad range of biomedical applications of the neuropeptides that they contain, conotoxins. Meanwhile anatomy in some other neogastropod lineages strongly suggests that they have evolved similar venoms independently of cone snails, nevertheless their venom composition remains unstudied. Here we focus on the most diversified of these lineages, the genus *Vexillum* (the family Costellariidae). We have generated comprehensive multi-specimen, multi-tissue RNA-Seq data sets for three *Vexillum* species, and supported our findings in two species by proteomic profiling. We show that venoms of *Vexillum* are dominated by highly diversified short cysteine-rich peptides that in many aspects are very similar to conotoxins. Vexitoxins possess the same precursor organization, display overlapping cysteine frameworks and share several common post-translational modifications with conotoxins. Some vexitoxins show detectable sequence similarity to conotoxins, and are predicted to adopt similar domain conformations, including a pharmacologically relevant inhibitory cysteine-know motif (ICK). The tubular gL of *Vexillum* is a notably more recent evolutionary novelty than the conoidean venom gland. Thus, we hypothesize lower divergence between the toxin genes, and their ‘somatic’ counterparts compared to that in conotoxins, and we find support for this hypothesis in the molecular evolution of the vexitoxin cluster V027. We use this example to discuss how future studies on vexitoxins can inform origin and evolution of conotoxins, and how they may help addressing standing questions in venom evolution.

## Introduction

The order Neogastropoda is a large and successful group of marine gastropod molluscs comprising over 18,000 described species (MolluscaBase). Most neogastropods are active predators or blood-suckers (Taylor et al. 1980), and many have developed unique biochemical innovations to assist hunting and defense (Olivera et al. 2014; Ponte & Modica 2017). The best known of them are venoms of *Conus* comprising structurally diverse oligopeptides, *conotoxins*, that cause devastating physiological effects in preys, and may be deadly for humans (Kohn 2018). Due to their ability to selectively block wide array of ion channels in the nervous system, conotoxins are one of the major highlights in the natural products based pharmacology (Prashanth et al. 2014; Safavi-Hemami et al. 2019). They are typically short cysteine-rich peptides, with a high proportion of post-translationally modified residues (Terlau & Olivera 2004). Conotoxin precursors have a uniform structure, comprising a signal sequence, a pro-region, and a mature peptide domain (Terlau & Olivera 2004; Puillandre et al. 2012). Whereas signal regions are typically highly conserved, the mature peptide domains evolve under strong positive selection, and were estimated to be among fastest evolving animal peptides (Chang & Duda 2012). Whereas cone snail venoms attract broad interdisciplinary interest, the fact remains barely acknowledged that venoms, likely similar to those in cone snails, are present in some other neogastropod lineages unrelated to Conoidea.

Conotoxins are synthesized in a specialized tubular venom gland, an evolutionary innovation of the hyperdiverse superfamily Conoidea (Puillandre et al. 2016; Abdelkrim et al. 2018), a homologue of the commonly found in neogastropods mid-gut gland of Leiblein, gL (Ponder 1973; Kantor 2002). Typically, gL has a spongy structure, and the use of its secretion for envenomation is unlikely: the duct of gL opens into the mid-oesophagus behind a distinctive valve of Leiblein (vL), which prevents any particle or fluid transport from mid-oesophagus anteriorly (Kantor & Fedosov 2009). However, several unrelated neogastropodan lineages beside Conoidea have evolved a massive tubular compartment in their gland of Leiblein. Its acquisition was invariantly accompanied by a modification or a complete loss of vL (Ponder 1973; Kantor & Fedosov 2009; Fedosov et al. 2017), thus effectively setting the stage for venom production and delivery. Several lines of evidence suggest that each neogastropod lineage possessing such derived morphology uses venom to subdue and kill the prey (Maes & Raeihle 1975; Olivera et al. 2014; Fedosov et al. 2019).

In the present study, we focus on the most diversified of these lineages, the genus *Vexillum*. We demonstrate the existence of venom in two *Vexillum* species, based on a comprehensive transcriptomic analysis of two tissues, supported by proteomic profiling. We show that venoms of *Vexillum* are dominated by highly diversified short cysteine-rich peptides that we name *vexitoxins* that in many aspects are very similar to conotoxins. Vexitoxins possess the same precursor organization, display overlapping cysteine frameworks and share several common post-translational modifications with conotoxins. Some vexitoxins show detectable sequence similarity to conotoxins, and are predicted to adopt similar domain conformations, suggesting that they have the same or similar molecular targets. Furthermore, we show that multiple unrelated vexitoxins contain the inhibitor cystine knot (ICK) motif (Pallaghy et al. 1994), which is present in many pharmacologically relevant animal toxins, including the conotoxin-based prialt (Robinson & Norton 2014). Therefore, vexitoxins have significant potential to become a novel source of bioactive peptides for drug development and neuroscience research.

## Results

### General characterization of the transcriptome datasets

A total of thirteen transcriptomic datasets were generated for four species of *Vexillum* (Table 1). Two tissues, salivary gland (sg) and tubular gland of Leiblein (gL) were sequenced with two replicate specimens for the three species, *Vexillum coccineum* (Vc), *Vexillum vulpecula* (Vv) and *Vexillum melongena* (Vm). Additionally, a smaller species, *Vexillum crocatum* was sequenced as a single pooled sample, containing salivary glands and tubular glands of Leiblein of two specimens (Fig. S1). The generated datasets are similar in terms of the number of reads per sample and in read quality metrics. The obtained Trinity assemblies were comparable in the BUSCO completeness (consistently slightly lower in sg compared to the gL of the same specimen), and were slightly lower in *V. melongena*, compared to *V. coccineum* and *V. vulpecula*. The assembly quality of the latter two species is comparable to that in the comprehensively sequenced *Conus* datasets (Barghi et al. 2015; Abalde et al. 2018). Furthermore, the proteomic data was obtained for three individuals of each species, *V. coccineum* and *V. vulpecula* to support and expand the transcriptomic analysis. Therefore, we mainly focus on the putative venom components identified in these two species, but also discuss sequences obtained from *V. melongena* and *V. crocatum* where relevant.

**Table 1.** Analysed transcriptomic datasets

| Species | Spm | Dataset | Total reads | N contigs<br>after<br>clustering | BUSCO<br>Mollusk<br>dataset<br>(%) | BUSCO<br>Metazoa<br>dataset<br>(%) | Mapped to<br>clustered<br>assembly<br>(%) |
| --- | --- | --- | --- | --- | --- | --- | --- |
| *V. coccineum* | #8Vc | #8Vcsg | 31,422,354 | 236,084 | 27.3 | 53.8 | 82.4 |
| *V. coccineum* | #8Vc | #8VcgL | 34,686,405 | 290,275 | 34.9 | 64.7 | 78.5 |
| *V. coccineum* | #9Vc | #9Vcsg | 32,132,355 | 249,943 | 33.9 | 64.9 | 83.6 |
| *V. coccineum* | #9Vc | #9VcgL | 29,313,679 | 302,835 | 36.1 | 68.7 | 79.8 |
| *V. vulpecula* | #13Vv | #13Vvsg | 32,942,762 | 124,115 | 20.8 | 43.3 | 87.6 |
| *V. vulpecula* | #13Vv | #13VvgL | 37,148,759 | 212,056 | 31.7 | 60.6 | 81.8 |
| *V. vulpecula* | #14Vv | #14Vvsg | 31,836,954 | 103,025 | 17.2 | 37.5 | 88.7 |
| *V. vulpecula* | #14Vv | #14VvgL | 26,942,349 | 291,987 | 38.3 | 67.1 | 77.6 |
| *V. melongena* | #33Vm | #33Vmsg | 34,649,613 | 74,707 | 16.1 | 35.6 | 82.3 |
| *V. melongena* | #33Vm | #33Vmgl | 32,781,696 | 149,065 | 27.1 | 53.2 | 82.0 |
| *V. melongena* | #35Vm | #35Vmsg | 38,273,786 | 87,825 | 21.0 | 47.0 | 85.6 |
| *V. melongena* | #35Vm | #35Vmgl | 36,993,835 | 124,406 | 27.1 | 53.5 | 84.7 |
| *V. crocatum* | #44Vr | #44VrsggL | 34,535,756 | 154,716 | 32.2 | 60.7 | 80.8 |

The coding DNA sequences (CDSs) predicted from the assembled contigs were filtered to keep only the CDSs that encode secreted peptides – these start with a C-terminal signal sequence, but lack a transmembrane domains. A total of 73,945 such CDSs were predicted in four *Vexillum* species; they were pooled and clustered with two alternative approaches: i) based on the identity of the signal sequence, with PID 0.65 - (Lu et al. 2020), and ii) based on the orthogroup inference. In further analysis we focus on the highly expressed clusters of CDS, so we built a reduced data set. If any CDS of a signal sequence based cluster, or of an orthogroup showed a TPM value exceeding 200 in any of the specimens, all members of this cluster or orthogroup, were added to the reduced data set. Thus compiled reduced data set included 3,308 CDSs that were subjected to manual curation to re-classify them to a final set of clusters that would reflect CDS sequence similarity but avoiding cluster oversplitting. We only kept CDS clusters comprising two or more CDSs, so the final data set comprised 235 clusters with 2,187 CDSs. Of these, 850 and 817 CDSs represented putative venom components of *Vexillum coccineum* and *V. vulpecula* respectively.

### Proteome and peptidome analysis

The main goal of proteomic analysis was to generate support for the venom components predicted based on the transcriptomic data. Because a notable proportion of these putative toxins were predicted to be rather short peptides, and could be passed to mass-spectrometric analysis without a preceding digestion, for each tissue, we analyzed both, the peptidome obtained from the native low molecular weight peptide fraction) and the proteome, generated from the trypsin-digested longer proteins (> 10 kDa).

The peptidome samples were directly subjected to LS-MS/MS analysis following with *de novo* sequencing by the PEAKS software, while the mass-spectra obtained from the digested samples were searched against the databases derived from the tanscriptomic data using conventional proteomic approach (for more details see the Material and Methods section) – Table 2.

**Table 2.** Results of the proteomic analysis of 12 *Vexillum* samples.

| Dataset | Shotgun proteomics |  |  | PTMs included |  | De novo sequencing |  |  | Total<br>CDS |
| --- | --- | --- | --- | --- | --- | --- | --- | --- | --- |
|  | peptides | CD<br>S | Summed<br>peps./CD<br>S | carboxyE<br>peps./CD<br>S | HydroxyP<br>peps./CD<br>S | peptides | CDS | summed<br>peps./CD<br>S |  |
| #3VcgL | 510 | 321 |  |  |  | 190 | 168 |  |  |
| #4VcgL | 527 | 329 | 727/439 | 104/101 | 104/146 | 46 | 83 | 254 / 211 | 479 |
| #6VcgL | 402 | 298 |  |  |  | 56 | 71 |  |  |
| #3Vcsg | 318 | 178 |  |  |  | 23 | 54 |  |  |
| #4Vcsg | 477 | 291 | 581/374 | 51/58 | 81/101 | 74 | 98 | 180 / 122 | 390 |
| #6Vcsg | 291 | 225 |  |  |  | 129 | 112 |  |  |
| #10Vvg<br>L | 423 | 246 |  |  |  | 94 | 74 |  |  |
| #11Vvg<br>L | 509 | 296 | 619/371 | 63/70 | 109/125 | 55 | 50 | 137 / 84 | 399 |
| #p7Vvg<br>L | 350 | 239 |  |  |  | 43 | 48 |  |  |
| #10Vvsg | 419 | 235 |  |  |  | 158 | 143 |  |  |
| #11Vvsg | 436 | 244 | 543/322 | 25/46 | 60/93 | 91 | 119 | 262 / 175 | 349 |
| #p7Vvsg | 316 | 209 |  |  |  | 124 | 145 |  |  |

Among the four analyzed species-tissue series, the gL datasets generated slightly higher number of hits, and no outliers in the hits number were detected in any series. A largest number of 727 unique matches was obtained from the specimens of *V. coccineum* gL, and the lowest (543 matches) from *V. vulpecula* sg. These generated support for 439 and 322 CDS respectively, however a majority of these supported CDS correspond to non-unique matches, because most peptides have generated hits to multiple database entries, which we collectively refer to as as “protein group”. When carboxylated glutamic acid and hydroxy-prolyne were set as variable modifications, additional sets of peptides were matched, again with larger numbers of hits in the gL series, compared to the sg of the respective species. Finally, from 137 to 262 native peptides per tissue-species series were revealed by *de novo* peptide sequencing in the peptidome samples. The largest (479) and the smallest (349) total numbers of supported CDS corresponded to the series of gL of *V. coccineum*, and sg of *V. vulpecula* respectively. We calculated overlaps among samples within each series i) in the detected peptides derived from the trypsin-digested protein fraction (FigS2, top row), and ii) in the subsets of CDSs supported by these peptides (second row). Our results highlight a notable concordance among the analyzed replicates at the CDS level: from 44% to 70% of the supported CDSs, are supported by all three conspecific tissue replicates. Largest contribution to the proteomic support of the query CDSs was generated by the peptides detected with conventional database search from the trypsin-digested protein fraction, however, a sizeable contributions, were also made by the *de novo* protein sequencing, and with modified matching accounting for 2 wide spread in conotoxins PTMs (Fig. S2, bottom row). Subsequently, we aligned all the peptides detected from the matched masses to the matching query CDSs, and summed up the length of predicted mature peptide region of each CDS, supported by the detected peptides. This value was divided by the total length of the predicted mature region, and the resulting ratio used as a measure of support; we report it for three best supported CDSs of each putative toxins cluster inferred from the transcriptomic data. In 31 and 25 CDSs of *Vexillum coccineum* and *V. vulpecula* respectively, obtained proteomic data was also essential to correct predicted boundaries of the mature peptide region.

### Venom composition in Vexillum

Confident BLAST or HMMER hits were obtained for 309 and 294 CDSs of *V. coccineum* and *V. vulpecula* respectively, which constitute 36.4% and 36.0% of the putative venom components in these two species respectively (Fig. 2A). The transcripts with reference-based annotations belonged to 47 Pfam gene families. Proteins bearing shaker toxin (ShKT) domains and metalloproteases, mainly of astacin type were the most diversified of annotated clusters in both, the sg and the gL of both species (Figs 2B, C). Both, ShKT domain bearing proteins, and astacins showed high expression in both tissue types, with notably higher total expression levels of ShKT domain bearing proteins in sg (Fig. 2D). Other highly expressed classes of venom peptides included Prohormone-4, peptidases –S, –M, and –C, and CRISP, Lectin-C, hyaluronidases, chitinase (CBM_14), Abhydrolase, serine type protease inhibitors (ITIH3, Pacifastin, Serpin), von Willebrand domain bearing proteins.

**Figure 1.**
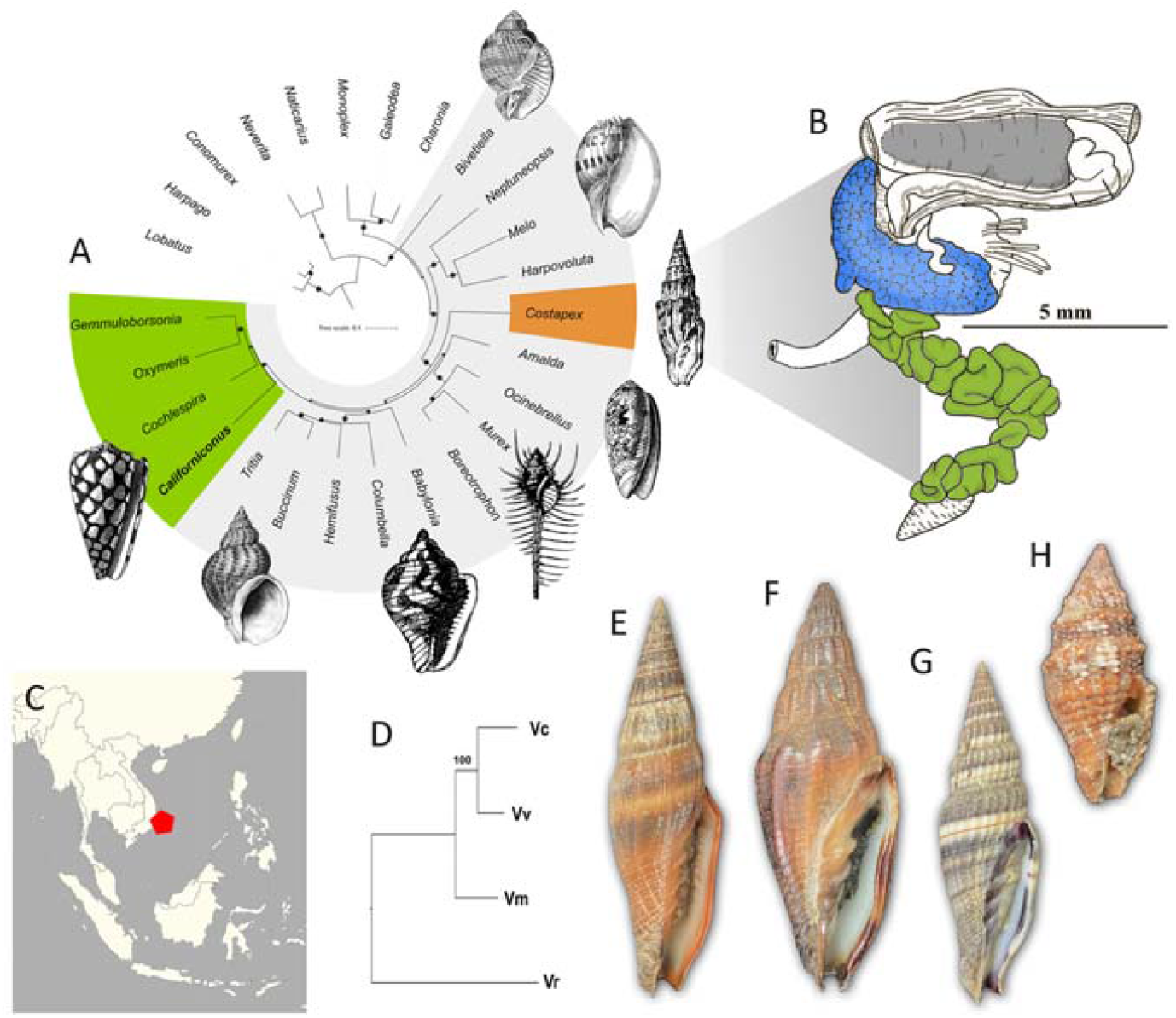
Phylogeny and morphology of *Vexillum*. A. Mitochondrial phylogeny of the Neogastropoda (after Uribe et al. 2021); the family Costellariidae represented by *Costapex baldwinae*; B. Foregut anatomy of *Vexillum vulpecula*, blue marks the salivary gland (**sg**), green – tubular gland of Leiblein (**gL**), grey - proboscis; C. Sampling location; D. Species tree of the four *Vexillum* species analyzed herein based on the ML analysis of concatenated aa sequences of 426 BUSCO loci 126,681 aa sites); E – H. Specimens dissected for transcriptomic analysis; E. *V. coccineum*; F. *V. vulpecula*; G. *V. melongena*; H. *V. crocatum*.

**Figure 2.**
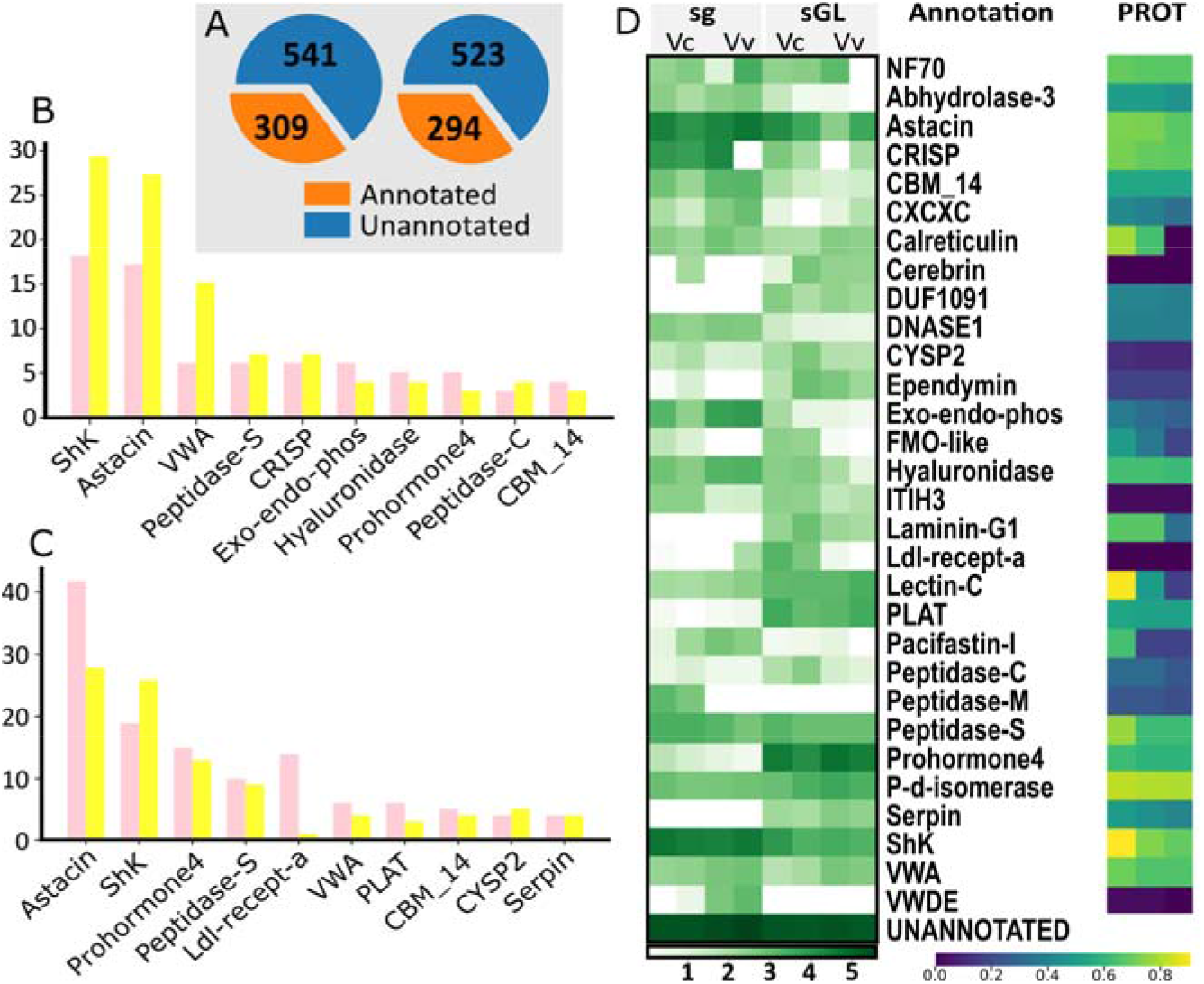
Major annotated clusters of transcripts in the sg and gL of *Vexillum* species. A. Proportions of annotated and unannotated transcripts in *Vexillum coccineum* (left) and *V. vulpecula* (right); B. Ten most diversified classes of annotated transcripts in salivary gland (sg), pink - *Vexillum coccineum*, yellow - *V. vulpecula*; C. Ten most diversified classes of annotated transcripts in gland of Leiblein; D. Heatmap of log10 transformed summed TPM expression levels of 30 most highly expressed annotated transcript classes per data set. On the right heatmap of the cluster support in proteomic data, where three cells in a horizontal row correspond to three CDS of a cluster best represented in our proteomic data. Color-coding corresponds to the proportion of the mature peptide length, represented in the proteomic data.

The small number of available replicates precluded statistically sound differential expression analysis; however, the contrasting expression levels in some venom components clusters can be noted (Fig. S3). For example, abhydrolase, CRISP, DNAse1, Exo-endo-phos, hyaluronidase, Pacifastin and ShKT bearing proteins show higher expression in sg (Figs 2D, S3). On the contrary, cerebrin, DUF1091, Ependymin, Laminin G1, Lectin C, PLAT-type metalloprotease, serpin and notably prohormone-4-like transcripts display higher expression in the gL.

While prohormone-4, lectins, CRISP, and hyaluronidases (as conohyal) have previously been identified in *Conus* venoms (Robinson et al. 2017; Fassio et al. 2019; Lebbe & Tytgat 2016), other *Vexillum* venom components are not typically reported from cone snails. Nevertheless, at least some of them: astacins, ShK-domain bearing proteins, peptidases, ab-hydrolases, serine-protease inhibitors are present in venom gland transcriptomes of the early-diverging cone snail lineages *Profundiconus* (Fassio et al. 2019), *Conasprella* and *Pygmaeconus* (Fedosov et al. 2021). Some of these transcripts typically show lower expression in cone snails, and were suggested to play an accessory role in envenomation, by facilitating spread of venom, or impairing the prey’s hemostasis (Fassio et al. 2019). The presence of these putative venom components in both the sg and gL of *Vexillum* as evidenced by both transcriptomic and proteomic data, suggests that secretions of both these glands play a role in envenomation. However, functional aspects of *Vexillum* venom components are still to be determined, and the priority here will be given to the putative toxins that we cover in further detail below.

### Proteins bearing ShKT domains are diversified and highly expressed in Vexillum

We identified a total of 98 complete transcripts of ShKT bearing proteins that can be classified to three gene superfamilies based on the identity of their signal sequence (Fig 3A). Because most predicted transcripts bear multiple ShK domains, we denote these clusters here as multiShKV1 – VexShKV3 (Gerdol et al. 2019). The members of these three gene superfamilies show major differences in both the numbers of ShKT-like domains that they comprise and the regions interleaving these domains. The only complete precursor of the small multiShKV1 gene superfamily, Vc00003648 is predicted to bear five ShKT-like domains (Fig. 3B). The three N-terminal domains show only limited identity to the canonical ShKT domain (HMMER evalue < E*10-2, and lack one or two cysteines). The transcripts of the large multiShKV2 gene superfamily encode up to five (e.g. Vv0001310, Fig. 3C), but typically three ShKT-like domains (e.g. Vc0000421, Fig. 3D). These transcripts feature a long low-complexity region with high proportion of charged (both positively and negatively) residues between the signal sequence and first ShKT-like domain. Finally, the majority of the multiShKV3 gene superfamily transcripts comprise only two ShKT-like domains (Figs 3E, F), and also contain a low-complexity region. This region spans up to 140 residues, and is composed of repeated short motif, starting with two negatively charged residues (typically DE), followed by 2-7 neutral residues. The log-10 transformed expression of the ShKT bearing proteins (Fig. 3A) shows transcripts’ expression in sg (blue) and gL (green), with the circular line marking a TPM 1000 expression (grey used for *V. crocatum* where the glands were pooled). It can be noted that multiShKV2 and multiShKV3 show contrasting expression patterns: the former is represented by about equal number of transcripts in sg and gL, but the highest expression transcripts are those in gL. Conversely, the multiShKV3 is dominating sg in both, the number of transcripts, and in their expression levels.

**Figure 3.**
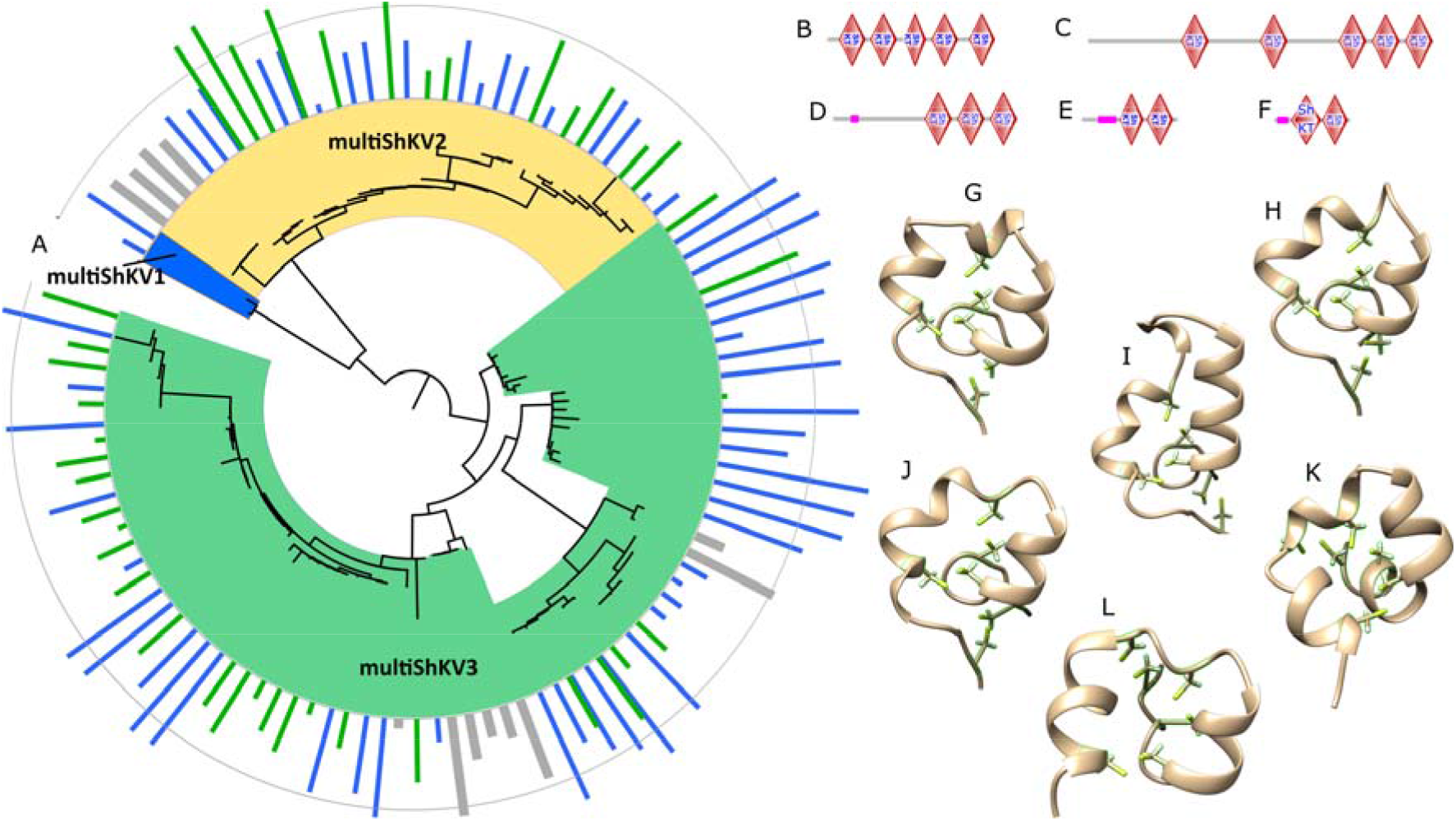
MultiShK proteins of *Vexillum*. A – E. Domain arrangement in five representative transcripts. A. Vc0003648 (multiShK1); B. Vv0001310 (multiShKV2); C. Vc0000412 (multiShKV2); D. Vc0000028 (multiShKV3); E. Vc0000358 (multiShKV3); F. Phylogenetic tree of the 98 identified complete multiShK protein precursors. The annotation corresponds to the log10 transformed TPM expression levels, shown in blue for sg, in green for gL, in grey – for polled tissues of *V. crocatum*. Outer circular line marks TPM expression level of 1000. G-L. Predicted 3D structures and inferred disulphide connectivity of the six structure types of *Vexillum* ShKT domains supported by proteomic data. G. Type I, Vc0000028, domain 1; H. Type II, Vc0000028, domain 2; I. Type III Vc0000358 domain 1; J. Type IV Vc0005635 domain 2; K. Type V Vv0001739 domain 2; L. Type VI Vc0005911 domain 2.

Previously identified ShK toxins of sea anemones are short neuropeptides, comprising six cysteine residues (Castañeda et al. 1995). They are potent potassium channel blockers, with high affinity to channels comprising a Kv1 subunit, especially of the Kv1.3 subtype (Pennington et al. 1995; Kalman et al. 1998). This makes them a valuable source of drug leads modulating immune functions: the Kv1.3 channels are crucial for terminally differentiated effector memory (TEM) T cells functioning, which are responsible for a wide range of autoimmune conditions. Many ShK toxins therefore have been chemically synthesized, and proved efficient in animal models of human autoimmune diseases (Chi et al. 2012; Tarcha et al. 2017). Of the total of 33 and 29 unique ShKT-like domains predicted in transcripts of *Vexillum coccineum* and *V. vulpecula* respectively, 17 and 12 respectively were supported by the proteomic data – all these are the domains encoded by the transcripts of multiShKV2 and multiShKV3 gene superfamilies. A total of 62 unique monoisotopic masses were detected in the proteomic datasets of *V. coccineum* that match the ShKT-like domain sequences, and a total of 31 unique masses support the *V. vulpecula* ShKT-like domains. We obtained high confidence 3D structure models (LTTD score typically above 90) for all the identified ShKT-like domains supported by the proteomic data (Figure S4). They demonstrated a high diversity of conformations that we classify into six general structural types, referred to as types I-VI (shown in the figures 3 G–L in the order of decreasing expression levels of the respective transcripts). The most common structure types I and II (Figs 3G, H) are encoded by both the multiShKV2 and multiShKV3 transcripts, and geometrically the closest match of both is the shaker toxin k of *Stichodactyla helianthus* (Figure S4). The structure types III, IV and V (Figs 3I-K), although share general features of ShKT domains, geometrically show higher resemblance to the pseudecins – the CRISP class toxins of Elapidae snakes targeting cyclic nucleotide-gated ion channels (Suzuki et al. 2008). Finally, one ShKT-like domain detected in *V. vulpecula* transcript V0001739 (Fig 3L) shows high structure resemblance to the natrin, a potent blocker of calcium-activated potassium (BK(Ca)) channels (Wang et al. 2005).

Although we do not have any direct evidence of the physiological activity of *Vexillum* ShK-like peptides, our data points at ion channels as their tentative targets. Indeed, the presence and the remarkable diversity of ShKT bearing proteins is predicted by the transcriptomic data of multiple species and specimens, and is further corroborated by the mass-spectrometric analysis. The very high expression of these transcripts in the secretory foregut glands suggests their significant role in the context of functionality of salivary glands and of the gland of Leiblein – i.e. presumably in envenomation. Finally, the detected sequence similarity of the *Vexillum* ShKT-like peptides with the sea anemone ShK toxins, and with the ion channel blockers of snake venoms potentially suggest that the ShKT-like peptides of *Vexillum* share same range of targets.

### Unannotated clusters of transcripts

The majority of the predicted secreted CDSs did not display any sequence similarity to the entries in the reference databases. Here we consider them together with the total of 32 CDSs that showed structure similarity with conotoxins (of these ten in *V. cocineum* and seven in *V. vulpecula*). The reason for it is that a large set of unannotated CDSs appears to share characteristic features of conotoxins, therefore the entire diversity of putative conotoxin-like transcripts is analyzed in the context of this similarity. These features are: i) the canonical precursor structure with a conserved signal sequence, and a rather short, variable mature domain represented by a single copy; ii) high number of cysteine residues in the mature domain that form distinctive cys-frameworks; iii) high number of post-translationally modified residues in the mature region.

The entire diversity of 1,580 unannotated secreted CDSs was classified into 146 clusters based on both, the identity of their signal sequences and the orthogroup inference; each cluster was assigned a digital code based on its summed expression (Table S1), supplemented by letters to reflect i) length of its constituent CDSs, and ii) degree of their sequence conservation. In total, 117 of these clusters demonstrated high expression in at least one of the profiled specimens (TPM ≥ 1000), or moderately high expression (TPM ≥ 100) across several specimens. In Figure 4A, the horizontal rows of cells that summarize cluster expression, are arranged based on the length of CDSs included in a cluster, and degree of sequence variation within a cluster (see colored ranges on the left). The clusters V001, V002, and V004 showed highest sequence diversity and extremely high expression in all analyzed datasets (Table S1). All three clusters appeared very heterogeneous. Despite the fact that sequences in each of them share a conserved signal sequence and recognizable sequence motifs in pre-and mature regions, each cluster included several distinctive major orthogroups (Figs S5-S7). In general, each of these three clusters showed notably higher expression levels in sg compared to gL (Fig 4A). Otherwise, it can be noted from the figure 4A that the clusters of medium-sized CDSs (entire precursor longer than 40 aa, but shorter than 200 aa), with over 10% variable aa sites (blue bar on the left) are much broader represented in gL than in sg.

**Figure 4.**
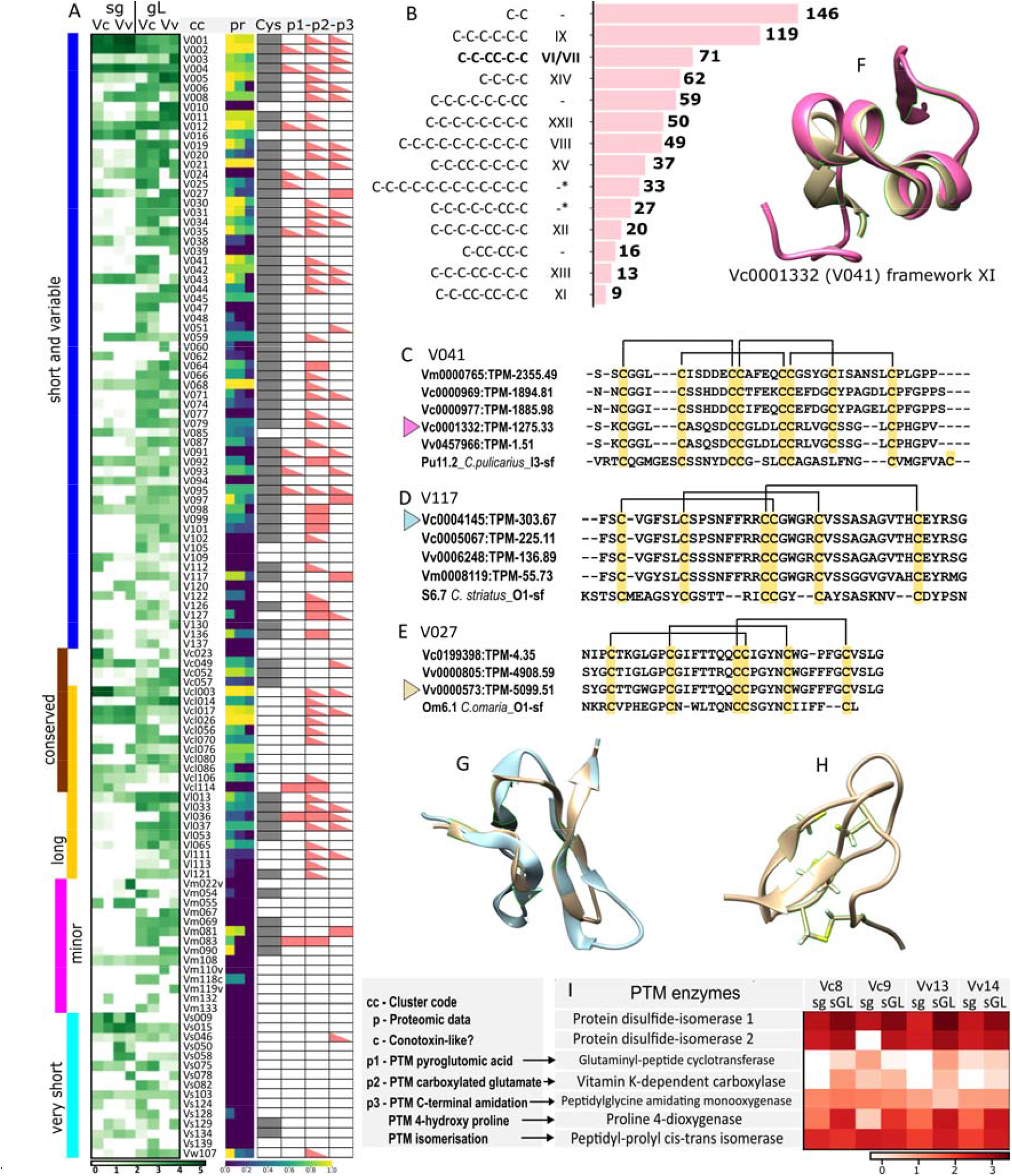
Expression and structural features of the unannotated *Vexillum* transcript clusters. A. heatmap of log 10 transformed expression of 118 unannotated clusters of transcripts in sg and gL transcriptomes of *Vexillum coccineum* and *V. vulpecula*. Column pr – support of clusters in proteomic data (markup like in Figure 2). Column c: grey marks presence of a conserved cys-framework across the sequences of a cluster, or of several compatible frameworks. Columns p1 – p3 – prediction of three PTMs most commonly found in conotoxins: p1 – N-terminal pyroglutamic acid, p2 – carboxy-glutamate, p3 – C-terminal amidation. B. Most common Cys-frameworks in unannotated clusters of putative *Vexillum* toxins. C – E. Mature peptide alignment in three clusters of vexitoxins with closest conotoxin matches. C. Cluster V041. D. V117. E. V027. F. Superposition of the vexitoxin Vc0001332 VS *Conus tulipa* conotoxin ρ-conotoxin TIA. G. Superposition of the vexitoxin Vc0004145 VS *Conus geographus* conotoxin GS. H. 3D structure of the vexitoxin Vv0000573. I. Heatmap of log 10-transformed expression level of seven key PTM enzymes in analyzed transcritomes of *Vexillum coccineum and V. vulpecula*.

**Figure 5.**
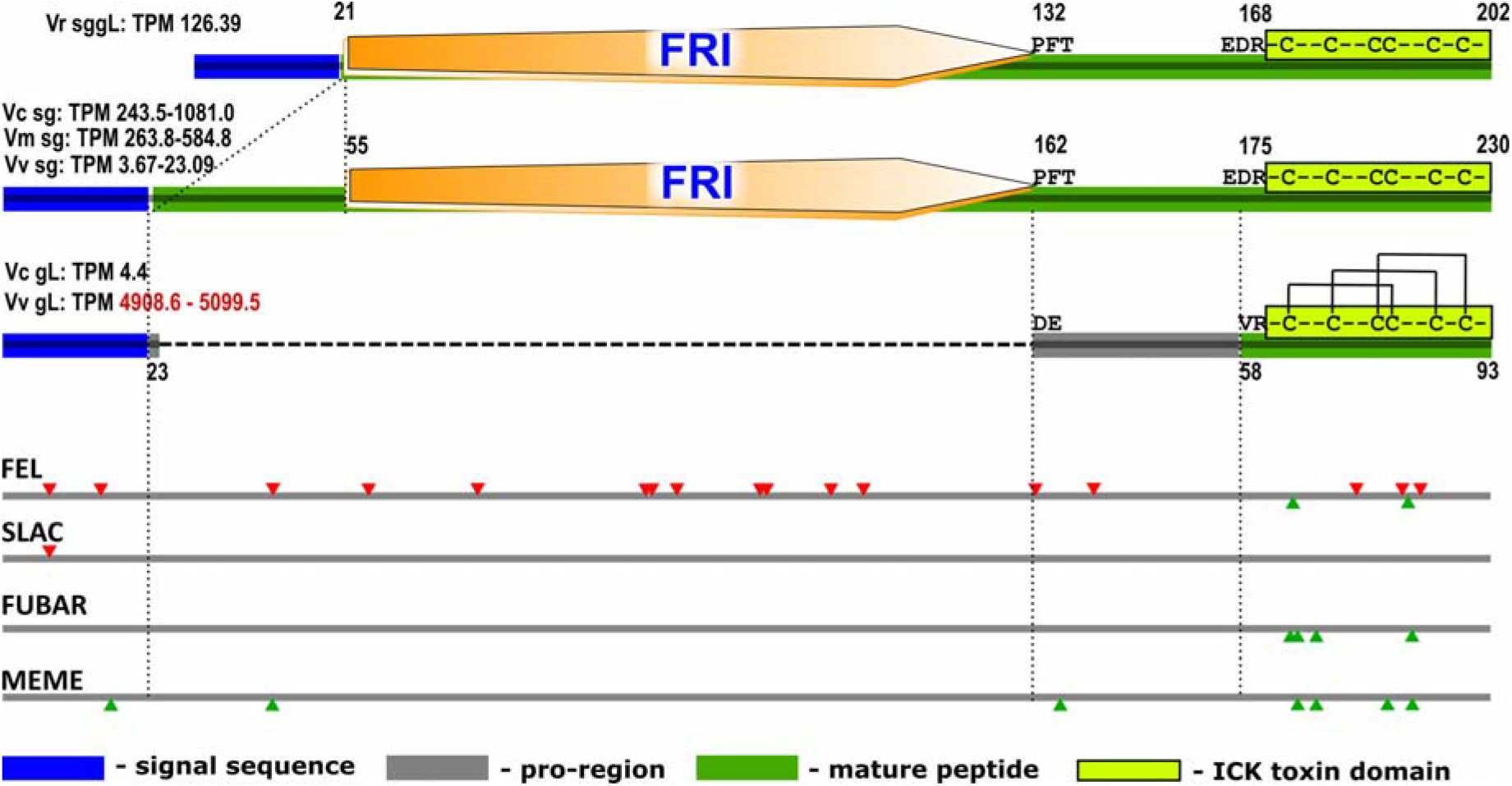
Precursor structure and evolution of the cluster V027 sequences in *Vexillum*. A. Precursor structures in V027. Top - long orthogroup, Vr0003450; Middle – long orthogroup, Vc0001640; Bottom – short orthogroup Vv0000573. B. Codons under negative selection (red), pervasive positive selection, identified by FEL, FUBAR, or SLAC, or episodic diversifying selection (identified by MEME) (green).

In the column ‘c’ of Figure 4A, we highlighted in grey those clusters, where mature regions of complete CDSs comprise at least two cysteine residues, and share the same or display compatible Cys-frameworks across each cluster (except V001, V002, and V004, where some variation was permitted). Fifty-five clusters can be considered as sharing structural features of conotoxins: they comprise cys-rich precursors whose length matches the length range of conotoxins. Of a total 942 complete transcripts in these clusters, 445 (or almost half) encode mature toxins with canonical Cys frameworks known from conotoxins. Of the 14 most common frameworks that are shared by no less than 10 predicted CDSs, nine are canonical frameworks known in conotoxins (Fig. 4B). For example, the framework IX found in 119 *Vexillum* CDSs is present in most P-superfamily conotoxins (Fedosov et al. 2012; Robinson et al. 2014), and the framework VI/VII, known also as the inhibitor cysteine knot (Robinson & Norton 2014; Lavergne et al. 2015), is most common in the O-, H- and N-conotoxin superfamilies. Two further frameworks marked with an asterisk are rather exotic for conotoxins (Lavergne et al. 2015). The VI/VII framework shared by 71 identified putative toxins of *Vexillum* is the third most common in our data set. The O1-superfamily conotoxins with the framework IV/IIV are potent blockers of voltage gated ion channels targeting Na^+^ channels (pharmacological families δ-, and μ-), K^+^ channels (κ-), and Ca^2+^ channels (ω-), and therefore are of great interest for drug development (Robinson & Norton 2014; Safavi-Hemami et al. 2019). In particular, the first conotoxin approved by FDA for clinical use the ω-conotoxin MVIIA (Prialt) possesses this cys-framework. The remarkable sequence diversity of framework IV/IIV toxins in *Vexillum* may suggest a similar scope of their molecular targets, and if proved true, *Vexillum* toxins may become a rich source of neuropeptides of high relevance for biomedical research and drug development.

The mature toxin alignments of three clusters that have displayed detectable similarity to conotoxins are showed in Figures 4C-E, their disulphide connectivity was inferred from the reconstructed high confidence 3D models (Figs 4E-H, respectively). The CDS Vc0001332 (cluster V041) has a rather uncommon cys-framework XI with four disulfide bounds. While its predicted sequence is closest to that of *Conus pulicarius* I3-superfamily conotoxin Pu11.2 (Fig 4C), the core of the predicted structure shows highest similarity to the much shorter ρ-conotoxin TIA (A-superfamily) of the fish-hunting species *Conus tulipa* (Fig 4F). The putative *Vexillum* toxins Vc0004145 and Vv0000573 both contain a ICK motif with its signature connectivity 1-4, 2-5, 3-6 (Figs 4G, H), and show closest sequence similarity to the S6.7 of *Conus striatus*, and to Om6.1 of *Conus omaria* respectively (both O1-superfamily). The modeled 3D structure of the Vc0004145 showed a close match to that of the synthetic μ-conotoxin GS (Hu et al. 2012) of *Conus geographus* (Fig 4G). Finally, some longer *Vexillum* toxins, such as the Vv0000706 (cluster V064), and Vc0004790 (V136) contain 12 cysteins which are predicted to fold into two ICK-like structures. The structure search on the obtained PBD files detected their highest structure similarity to the cyriotoxin-1a of the spider *Cyriopagopus schioedtei* (Fig. S7).

To estimate, whether the predicted *Vexillum* toxins bear same PTMs as do conotoxins, we summarized the PTM predictions obtained from ConoPrec (Fig 4A, columns p1 – p3), and corroborated these by the expression data of the corresponding PTM enzymes in the sg and gL transcriptomes (Fig. 4I). Our results suggest that these PTMs are likely to be quite common in *Vexillum* toxins. Among the predicted PTMs, the gamma-carboxylated glutamate was most commonly predicted (395 putative toxins from 56 clusters), however, the reliability of this PTM prediction from the primary protein sequence is questionable (Shah & Khan 2020). When glutamate carboxylation was set as a variable modification to expand the search of MS data, we recovered 25 to 104 additional unique monoisotopic masses per tissue-species series, with larger number of additional hits in gL compared to conspecific sg samples (Table 2). This suggests that if not a most common PTM, gamma-carboxylated glutamate at least occurs with a detectable frequency. The N-terminal amidation was predicted as the second most common PTM (236 putative toxins from 32 clusters). Furthermore, we detected presence of the seven essential PTM enzymes in the analyzed transcriptomes (Fig 4I). Protein disulfide isomerase, prolyl 4-hydroxylase (P4H), and peptidyl-prolyl cis-trans isomerase (PPI) showed highest expression levels. There is a clear pattern with higher expression of all these enzymes in the gL compared to sg. Glutaminyl-peptide cyclotransferase (GPC), Vitamin K-dependent carboxylase (VKD), and peptidyl-glycine amidating monooxygenase (PAM), were detected in all gL transcriptomes, but the former two were lacking in two sg data sets. Nevertheless, there is no tissue-specific pattern in the expression of the latter three enzymes. The presence of these essential PTM enzymes in most analyzed transcriptomes supports the hypothesis that peptide products of sg, and particularly, gL feature same post-translational modifications as conotoxins.

### Cross-tissue recruitment exemplified by the V027 cluster evolution

A close inspection of the cluster V027 sequences revealed that they represent two orthogroups with considerable differences in sequence length and tissue expression specificity. The first orthogroup sequences are about 230 aa long and are expressed predominantly in salivary glands, with the expression levels varying among species. The second orthogroup sequences are 92-93 aa long, and are only detected in gL of *Vexillum coccineum* and *V. vulpecula* with a very low expression in the former, and conversely, a fairly high expression in the latter (TPM-∼5000). Both orthogroups share a high identity N-terminal signal sequence, and a short C-terminal 35 aa long fragment, with a ICK motif. The observed length difference between them is due to the presence in the first orthogroup sequences of a conserved 111 aa long region annotated as a ‘frizzled` domain by HMMER, and showing highest sequence similarity to the cys-rich domain of the FZD4 protein (Zhang et al. 2011). The first orthogroups sequences are predicted to cleave into two fragments, one corresponding to the N-terminal signal sequence, another one to a long C-terminal mature peptide combining the Fz-domain with its flanking regions, and the ICK-bearing domain. On the contrary, the second orthogroup sequences are predicted to be cleaved in a manner similar to conotoxin precursors: into three fragments, corresponding to i) a signal sequence, ii) a short pro-region, and iii) a short mature peptide, which exactly corresponds to the N-terminal ICK-bearing domain. This mature peptide sequence shows a detectable similarity to the omega-conotoxin Om6.1 of *Conus omaria* (Fig. 4E), and molecular modeling predicted it to adopt a conformation characteristic for omega toxins family (Fig. 4H). Three monoisotopic masses, uniquely matching the Frizzled domain aa sequence are detected in proteomic data on *V. coccineum*. Consistent with transcriptomic data, these peptides were present in all six analyzed samples (i.e. in both, the sg and gL), but were entirely lacking in *V. vulpecula*. On the contrary, peptides matching the V027 ICK motif were only detected in the gL samples of *Vexillum vulpecula* (#11VvgL, #p7VvgL), and were represented by two nearly identical peptides (16 aa and 17 aa long) that match the N-terminal half of the ICK of the transcript Vv0000805.

The reconstructed phylogeny of the V027 cluster sequences (Fig S8) and the orthogroups distribution across species suggest that the first (longer) orthogroup transcript structure is ancestral, and the one of the second orthogroup is derived. The selection analysis identified 17 sites of the precursor under the negative selection, and these sites are predominantly located in the Fz domain. Conversely, of eight sites identified across the alignment, subject to either pervasive positive selection (FEL, SNAP, FUBAR), or evolving under diversifying selection (MEME), five are within the ICK-bearing domain.

This pattern is consistent with the second orthogroup descending from the first one resulting from a gene duplication event that has occurred before the split of *V. coccineum* and *V. vulpecula*. Following the gene duplication, the second orthogroup sequences lost the Fz-domain, and acquired a cleavage site at the N-terminal boundary of the cys-rich region. Subsequently, the shortened mature peptide region was rapidly evolving under positive selection and gained high tissue-specific expression in gL of *Vexillum vulpecula*. The very high expression of the transcripts Vv0000573 and Vv0000805 in *V. vulpecula* gL, presence of the matching translation products in the proteome, and their 3D structure determined by the ICK, all point at the relevance of this cluster to envenomation. This example illustrates how the cross-tissue recruitment of a gene copy followed by its accelerated sequence evolution gives rise to a pharmacologically relevant venom component following predators’ speciation.

## Discussion

### Vexillum toxins a novel source of bioactive neuropeptides

The molecular targets of conotoxins – a wide array of ion channels, and receptors in nervous system and at neuro-muscular junction have made them promising source of analgesics and a potentially preferable treatment for long-term pain management (Safavi-Hemami et al. 2019). The relevance of conotoxins as pharmacological agents can be explained by the fact that venoms in some cone snail species were evolved specifically to subdue vertebrate preys (Olivera et al. 2014, 2015). In this perspective, the fish-hunting species of *Conus* (or more broadly, those venomous lineages that are specialized to hunt vertebrate preys), are the first priority for drug-discovery. While this logic formulates a ‘pragmatic’ approach to prioritizing targets of resource-consuming drug development process, it would lead to *a priori* elimination of many potentially valuable candidate molecules. For example, the α-conotoxin Rg1a acting as an inhibitor of the α9α10 nicotinic acetyl-choline receptors (nAChR), proved to be a potent analgesic (Bjørn-Yoshimoto et al. 2020; L et al. 2014), despite being produced by a worm-hunter species *Conus regius*. Similarly, sea anemones do not feed on vertebrate preys, nevertheless, ShK toxins have high affinity to the vertebrate subtypes of potassium channels (Pennington et al. 1995). These examples may be explained by either broad taxonomic distribution of relevant molecular targets, or by the existence of defensive components of venoms, which evolve to efficiently deter vertebrate predators, rather than to subdue a prey. The defensive venoms targeted to vertebrates may have a much broader distribution across animal lineages, compared to the predatory toxins targeted to vertebrates. Furthermore, the ancestral defensive venom compounds are believed have become the substrate for the evolution of a novel predatory toxin set enabling piscivory in *Conus* (Imperial et al. 2007). In this context, a broader sampling of venomous animal taxa is crucial to systematically explore their molecular adaptations to hunting and, as well, to defense, and to efficiently reveal novel bioactive compounds of potential interest for pharmacology.

In the present study, we make a first step towards documenting venom composition of a highly diversified, yet previously unexplored lineage of venomous gastropods, the genus *Vexillum*. Due to the small size of its glands compared to a venom gland of cone snails, collection of sufficient material for bioassays of *Vexillum* venoms is a challenging task. To overcome this challenge, we used RNA-Seq and shotgun proteomics approaches that both require little material to generate a high-quality multi-tissue, multi-specimen, and multi-species data set to enable a rigorous analysis of *Vexillum* venom composition. We uncover highly diversified short secreted peptides referable to CRISP neuropeptides class in both, the salivary gland, and specialized tubular gland of Leiblein of *Vexillum*. One distinct group of these neuropeptides are the shaker-like toxins. These are synthesized as multiShK proteins that constitute two highly diversified unrelated multigene families with contrasting expression patterns in sg and gL of *Vexillum*. The ShKT domains encoded by these proteins share crucial structural features of the sea anemones ShK toxins, and therefore are likely to share similar or related molecular targets. Because ShK toxins of sea anemones have proved efficient in treatment of some autoimmune conditions (Chi et al. 2012), due to their high affinity to Kv1.3 potassium channels, *Vexillum* counterparts of the ShKTs represent an interesting group of short peptides for further pharmacological characterization.

We analyze the vast diversity of short Cys-rich secreted peptides of *Vexillum*, for which no reference-based annotation could be retrieved, in the context of their similarity with conotoxins. In total 55 transcript clusters (47 supported by proteomic data), show structural features characteristic of animal toxins (in particular, of conotoxins): short mature domain, largely conserved Cys-framework, and the presence of some signature PTMs. Of them 141 and 127 complete transcripts were identified in *Vexillum coccineum* and *V. vulpecula* respectively that share canonical Cys-frameworks of known classes of conotoxins. These numbers fall well in the range of the per-species conotoxin diversity assessed from the venom gland transcriptomes of *Conus* (Fassio et al. 2019). Although we do not have functional data to support the claim that these predicted transcripts indeed encode potent toxins, we present strong evidence that i) their translation products do exist in the protein fraction of analysed secretory glands, and ii) structural features strongly suggest that at least a sizeable fraction of them are toxins. It is thus logical to propose that they target same physiological circuits of preys and predators as do the conotoxins. Therefore, by this study we establish a solid background for the subsequent functional characterization of identified *Vexillum* toxins.

### Comparative framework for venom evolution inference in Neogastropoda

Venoms have evolved over hundred times in independent metazoan lineages (Schendel et al. 2019), offering a unique opportunity for studying genetic underpinnings of repeated key traits apparition (Casewell et al. 2013; Zancolli & Casewell 2020). Being a key adaptation for predation and defense, venoms to a great extent affect species fitness and biology (Dutertre et al. 2014; Casewell et al. 2017). Setting up venom production requires novel specialized tissues and glands, in which a set of genes originally not related to the venomous function is recruited and modified to encode potent toxins. Most animal toxins represent rather few broad classes of proteins, such as cysteine rich secretory peptides (CRISPs), hyaluronidases, kunitz-phospholipase and serine-type proteases (Zancolli & Casewell 2020), but being broadly distributed across unrelated venomous animal taxa, they have been recruited from very different genomic backgrounds (Barua & Mikheyev 2021). This general trend to convergent evolution provides a unique opportunity to disentangle the interplay of lineage-specific and conserved mechanisms that govern recruitment and evolution of venom peptides. To enable such inference globally, a scalable comparative framework should be generated to cover entire phylogenetic diversity of venomous animals. Notwithstanding, taxonomically restricted fragments of such framework may yield deep insights into genomic underpinnings of evolution and regulation of venomous function. Currently, most efforts to this end focus on the well characterized taxa of venomous animals, mainly on snakes (e.g. Barua & Mikheyev 2019, 2021), and extending such studies to new system(s) will greatly magnify the power of comparative analysis. Essentially, such system can be seeded by a pair of distantly related taxa that have independently acquired venom function, and cone snails and *Vexillum* representing unrelated evolutionary successful radiations of venomous neogastropods are thus a perfect system.

Evolutionary histories and distributions of *Conus* and *Vexillum* display multiple parallels. Similar to *Conus, Vexillum* is species rich (encompassing about 390 species), and forms a crown group of its respective family, the Costellariidae (Kohn 1990; Fedosov et al. 2017). Similar to *Conus, Vexillum* underwent a major diversification in Miocene, and its present day diversity is mainly associated with tropical shallow waters of Indo-Pacific. Therefore, the adaptive radiations of *Conus* and *Vexillum* were likely shaped by the same set of factors, and acquisition of venom likely have played a major role in the success of both these taxa. Within this system, repeated recruitments of a novel specialized secretory tissue of gL allows comparative analysis of the genome evolution processes underpinning emergence of venom gene superfamilies, and establishment of their regulatory pathways. Because tubular gL has the same developmental origin in *Vexillum* and *Conus* (as stripped off dorsal oesophagus wall), the gene expression patterns in the ancestral tissues were presumably closely comparable among them. Conversely, sg is homologous **and** morphologically conserved across Neogastropoda, and also produces some classes of bioactive compounds in both cone snails and *Vexillum* (Ponte & Modica 2017; Biggs et al. 2008, the present study). This two-tissue system enables a comparative analysis of modes and tempos of molecular evolution, as well, as investigation of cross-tissue gene superfamilies recruitment between sg and gL. In the present study, we demonstrate an example of the sg-gL cross-tissue recruitment in the *Vexillum* V027 cluster.

After an initial gene duplication, the gene structure of the new paralog was modified to produce a short ICK-bearing toxin. Subsequently, after the divergence of *Vexillum coccineum* and *V. vulpecula*, the toxin sequence evolved under accelerated positive selection and gained high expression in the gL of the latter species. Interestingly, the mounting expression of this toxin in gL of *V. vulpecula* was accompanied by the reduction of the ancestral (long) paralog expression in the sg, suggesting that the functionality of their gene products may to some extent overlap. What we find remarkable in this example is that we were still able to capture the low expression counterpart of the ancestral (long) orthogroup in the sg of *V. vulpecula*. Furthermore, we detected a low-expression ‘prototype’ of the ICK-bearing toxin gene (the short orthgroup) in the gL of *V. coccineum*, where it is expressed alongside the ancestral orthogroup, but with an order of magnitude lower expression level. It is likely that these low-expression counterparts are not functional in the context of the biology of the respective species, and their observed expression is residual, and would have completely vanish if the divergence between *V. coccineum* and *V. vulpecula* was less recent.

The observed distribution of orthogroups across tissues and species of *Vexillum*, as well, as the distribution of ShKT-bearing transcripts, imply that there remains some functional overlap between the sg and gL in *Vexillum*, in relation to envenomation. If true, such overlap may generate a ‘highway’ for cross-tissue recruitment of venom components in the evolutionary young gL (Fedosov & Kantor 2010; Fedosov et al. 2017) by means of subfunctionalization (Hargreaves et al. 2014). Therefore, a sizeable fraction of venom components in *Vexillum* is likely to result from recent recruitment events, and so, despite the inherent quick divergence from the ancestral state, the structural or sequence similarity of these venom genes with their non-venomous paralogs may still be traceable. If this is true, *Vexillum* venoms provide an ideal system to study origin and early evolution of venomous function in general. Furthermore, outcomes of this analysis have a great potential to inform the evolution of conotoxins. Genomic source of conotoxin genes origin remains unknown, mainly because these genes evolve too fast, and the venomous function has originated in the ancestors of cone snails too long ago (Abdelkrim et al. 2018) for detection of the toxin genes ancestry to be possible. However, because *Vexillum* and *Conus* share a common ancestor within the Neogastropoda, their genomic background is largely the same. Therefore, venom evolution inference in *Vexillum* will give a shortcut to identifying the set of ancestral neogastropod genes amenable for venom function, and this knowledge, in turn, will generate sensible hypotheses on the evolutionary origin of conotoxins.

## Supporting information

Kuznetsova et al. supplementary figures 1-9

## Acknowledgments

We are grateful to the staff of joined Russian-Vietnamese tropical center for supporting sampling in Nha-Trang Bay. We thank Dr. Manolo Tenorio (University of Cadiz) for help with running AlphaFold, and Dr. Helena Safavi-Hemami (University of Copenhagen) for her comments on the manuscript, and Dr. Yuri Kantor (IEE RAN) for valuable discussion. The present research was supported by the RSF grant 19-74-10020 to AF.

## Data availability

The transcriptomic sequencing data are deposited in the NCBI SRA database, under the Bioproject PRJNA797643. Sequences of the predicted *Vexillum* toxins are provided in the supplementary data files. The essential Python scripts used for the data analysis are available at https://github.com/SashaFedosov/Vexillum/).

## Material and methods

### Specimen collection and tissue sampling

Specimens of four *Vexillum* species were collected by SCUBA diving in Nha-Trang Bay, Central Vietnam in May 2021. Five specimens of *V. coccineum* measuring 54 - 58.5 mm, five specimens of *V. vulpecula* (60 – 67.5 mm), and two specimens of *V. melongena* (51.5 and 52.8 mm) were collected at depths 5-8 meters in Dam Bay (Tre Island) on silted sand. Two specimens of *V. crocatum* (22.3 and 28.4 mm), were collected in a crevice of a vertical reef wall at depth 12 meters off Noi Island. All specimens were delivered in the onshore laboratory and kept in tanks with aeration overnight; dissections were performed on the following day. Two specimens of each species were dissected for transcriptomic analysis, and three additional specimens were dissected for each, *V. coccineum* and *V. vulpecula* for proteomic analysis. Prior to the dissection, each specimen was photographed, then a vise was used to destroy the shell, and the body was promptly dissected to excise the salivary gland (sg) and the tubular gland of Leiblein (gL). These were preserved individually, for each specimen except *Vexillum crocatum* – for the latter species two sg and two gL were pooled in a single sample (44VrsggL). Tissues for transcriptomic analysis were preserved in RNA*later* (ThermoFisher), kept 24 hours at room temperature, and then stored at −20°C until dissection. Samples for proteomic analysis were immediately frozen in liquid nitrogen, and kept at −70°C until further processing. A fragment of foot was clipped from each dissected specimen and preserved in 95% ethanol to confirm species identity by means of DNA-barcoding.

### RNA Isolation, and sequencing

RNA was extracted from sg and gL tissues of *Vexillum* using the standard Trizol method. Bioanalyzer traces were used to assess total RNA quality and determine suitability for sequencing. The cDNA libraries for Illumina pair-end sequencing were then prepared following the automated polyA RNAseq library prep protocol. All libraries were sequenced on the Illumina HiSeq 4000 patform, at the sequencing facility ‘Genoanalitica’ (*V. coccineum* and *V. vulpecula*), or at the genomics core facility of Skolkovo Institute of Science and Technology (*V. melongena* and *V. crocatum*).

### Transcriptome assembly and reference based annotation

The raw reads were quality checked using FastQC, and then filtered to remove putative contamination by running FastQ-Screen v0.14.1 (Wingett & Andrews 2018), with Bowtie2 (Langmead & Salzberg 2012) mapper. The reads were mapped to 26 genomes, including those of Human, mouse, yeast, *Drosophila, Arabidopsis, E. coli* and *Cutibactrium acnes*, as well, as to the genomes of the other organisms that were library-prepped, or sequenced alongside our *Vexillum* samples. The reads that did not map to any genome were retained for assembly. They were trimmed using Trimmomatic v0.36 (Bolger et al. 2014) with the following parameters: ILLUMINACLIP option enabled, seed mismatch threshold = 2, palindrome clip threshold = 40, simple clip threshold of 20; SLIDING WINDOW option enabled, window size = 4, quality threshold = 15; MINLEN = 36; LEADING = 3; TRAILING = 3and assembled using Trinity v2.11 (Grabherr et al. 2011) with default parameters (kmer size=25, transcript identity=0.98, minimal contig length=200. We used RSEM v1.3.1 (Li & Dewey 2011) with the Bowtie2 mapper, to produce TPM-based measures of transcript abundances, according to the most common practice (Phuong et al. 2016; Abalde et al. 2018; Fedosov et al. 2021). We did not perform TMM correction among samples because it requires generating single assembly for each species which we abandoned, because it resulted in a reduced number of reads mapped), and still does not allow for normalization among species. Same Trinity assemblies were used to evaluate completeness of the datasets based on two BUSCO datasets, the metazoan dataset (954 loci), and the Mollusca dataset (5295 loci) (Waterhouse et al. 2018). We retrieved TPM expressions levels for the complete BUSCOs extracted from each dataset, and ranged them by increasing TPM expression level. We arbitrarily denoted the TPM expression level corresponding to the 25 percentile of this distribution as the minimal confidence threshold: the predicted transcripts of this dataset with lower expression levels were discarded.

Coding DNA sequences (CDSs) were predicted from the Trinity assembly using ORFfinder (NCBI), keeping only those CDSs that comprised over 35 amino acid residues. First, they were further filtered to remove possible cross-contaminations by applying the following filter: if an CDS showed TPM expression level ≤ 0.01 relative to an identical CDS from some other specimen sequenced at the same facility, the former CDS was removed from the dataset (custom Python script PS1.py). Then, a non-redundant catalog of all remaining CDSs was built for each species, where CDSs were ranked by their TPM expression levels summed across specimens. The secreted gene products were identified as CDSs that contain a signal sequence, identified by SignalP v5.0 (Nielsen 2017) with a D-value, D>≥0.7, but lack a transmembrane domain, detected by phobius v1.01 (Käll et al. 2007). To detect putative assembly errors, in the CDSs that passed this filter, we retrieved the per-base coverage data for each CDS using samtools_depth function, and a custom Python script PS2.py, and checked it to ensure that there are no abrupt shifts in the transcript coverage. The subset of CDSs that passed these filters was subjected to a sequence-based annotation by means of BLASTp against the manually curated SWISS-Prot database (Bairoch & Apweiler 2000), and the structure-based annotation using HMMER v3.2.1 (Finn et al. 2011) against the database of Hidden Markov Models, HMMs derived from PFam (Mistry et al. 2021).

### Transcripts de novo annotation

Because there are no genomic resources available for *Vexillum* or any closely related lineage of Neogastropoda, we expect that only a subset of *Vexillum* venom components can be revealed by the reference based annotation (Fedosov et al. 2021). Therefore, we first performed CDSs clustering, and then those clusters that showed high expression in either sg or gL were annotated. First, we combined four CDSs catalogs corresponding to the secreted gene products of our four species in single file, and then clustered them using two alternative approaches. Because signal sequence is highly conserved in *Conus* toxins, the classification of conotoxin gene superfamilies relies on its identity (Puillandre et al. 2012), and so a conotoxin can be assigned to a gene superfamily based on the signal sequence matching. Most conotoxin gene superfamilies show 0.55 – 0.7 Percent identity (PID) of the signal sequence (Kaas et al. 2012). We used CD-Hit (Fu et al. 2012) with two values of PID, 0.6 and 0.65 to generate two alternative sets of clusters for our ORFs. However, the algorithm of CD-Hit tends to neglect similarity of longer sequences, and therefore, they may end up in different clusters despite sharing a highly identical region. As an alternative approach, we inferred orthogroups based on the whole CDS comparisons by Orthofinder2 (Emms & Kelly 2019). Because subsequent annotation required laborious manual curation, we only focused on the highly expressed transcript clusters. We built a reduced dataset, which contained all sequence of a given SS-cluster, or of a given orthogroup if at least one of the CDSs in this orthogroup/SS-cluster had a TPM expression value exceeding 200 (custom Python script PS3.py). This reduced dataset included 3308 CDSs, representing 1056 orthogroups and 623 signal sequence based clusters (PID=0.65). This dataset was manually curated to establish optimal cluster breakdown based on i) signal sequence identity, ii) orthogroup inference and iii) available reference-based annotation. When several alternative cluster breakdowns were suggested, whole precursor alignments were built and examined to identify the best breakdown. After the removal of truncated CDSs and orthogroups / SS-clusters of single-CDS, the final dataset included 2187 CDSs allocated to 235 putative toxins clusters. Further annotation was performed for the 146 clusters comprising 1,580 CDSs (of them 1341 complete) that either were classified as conotoxins by HMMER, or had not returned any hits in the reference based annotation.

Each cluster was assigned a code based on its summed expression, length of included CDSs, and degree of their sequence conservation. We arranged all 146 clusters based on the summed expression in descending order and assigned each cluster a number based on its position in this ranking (Table S1). Then we considered separately the 14 clusters comprising very short CDSs (<40aa), clusters with CDSs of intermediate length (exceeding 40aa, but less than 200 aa), and long CDSs (length exceeding 200 aa). The letters ‘s’ and ‘l’ were appended to the cluster code for clusters of ‘short’ and ‘long’ CDSs respectively. Letter ‘c’ was appended to the codes of those clusters that showed high level of CDS sequence conservation (length variation < 1% of the average complete CDS length, and proportion of variable sites in the aa alignment <10%). Finally, a few clusters that contained only 2-3 CDSs, were denoted as ‘minor’, and letter ‘m’ was appended to their codes. Regardless of the expression, length, and degree of conservation, these 146 clusters were treated as putative toxins, and their precursor structure was analysed by Conoprec (available at http://www.conoserver.org/index.php?page=conoprec), to establish the domain breakdown, and to identify putative post-translational modifications (PTMs) and canonical cys-frameworks.

To generate additional support for the PTMs identified by ConoPrec, we performed a search for respective PTMs enzymes in sg and gL of *V. coccineum*, and *V. vulpecula*. We accessed all Uniprot sequences of the following enzymes identified in caenogastropod mollusks: Glutamynil-peptide cyclotransferase (PTM: N-terminal pyroglutamic acid), Vitamin K-dependent carboxylase (PTM: *γ*-carboxylated glutamic acid), peptydylglicine amidating monooxygenase (PTM: C-terminal amidation), and Prolyl 4-hydrolase, carrying out PTM of proline to 4-hydroxy proline, the fourth common PTM in conotoxins. Furthermore, we accessed Uniprot sequences of the two enzymes that have been shown to play important role in folding of conotoxins: protein disulfide isomerase and peptidylprolyl cys-trans isomerase (Safavi-Hemami et al. 2010). All predicted transcripts in the Trinity assemblies of *V. coccineum* and *V. vulpecula* that generated a blastx hit to any of the PTM enzymes sequences with aligned length ≥ 50% of the respective database entry length, and the BLAST e-value ≤ −25 were recorded, and their expression levels were summed up.

We predicted 3D structure of mature peptide domains for a few putative toxins for which we recovered sufficient support in the proteomic data. The structure modeling was performed in the ColabFold notebook (available at https://colab.research.google.com/github/sokrypton/ColabFold/blob/main/beta/AlphaFold2_advanced_beta.ipynb), which implements the recently released AlphaFold2 (Jumper et al. 2021). The multiple sequence alignments were built using the MMseqs2 algorithm, following by the prediction of five best spatial models, based on their mean pLDDT (Local Distance Difference Test) scores. The model with the highest LDDT-score was refined using Amber (Case et al. 2005)(available as a part of the ColabFold workflow), and then visualized in Chimera v1.15 (Pettersen et al. 2004) to infer cysteine connectivity. This model was also used for the structure-based search, which we performed using the online RUPEE protein structure search (Ayoub & Lee 2019) (available at https://ayoubresearch.com/) against the SCOPe database (Chandonia et al. 2019).

### Phylogenetic inference

To reconstruct species tree of the four analyzed *Vexillum* species, the complete BUSCOs extracted from each transcriptomic dataset were merged to build a non-redundant catalog of BUSCOs for each species. The amino acid sequences were aligned separately for each BUSCO locus using MAFFT v7.407 (Katoh & Standley 2013), and then a concatenated matrix was built from those 426 BUSCO loci from the Mollusca dataset that were present in all four *Vexillum* species (custom Python script PS4.py). This matrix comprising a total of 126,681 aligned aa sites was passed to IQtree v1.6.9 (Nguyen et al. 2015) as a single partition for phylogenetic inference.

To reconstruct gene tree of the V027 vexitoxin cluster, the nucleotide sequences of the ten complete CDSs identified in this cluster were codon-aligned using MACSE v2 (Ranwez et al. 2018). Then we ran IQtree with 1000 ultra-fast bootstrap iterations, treating three codon positions in the alignment as three separate partitions.

### Evolutionary analysis

The codon-aligned coding sequences of *Vexillum* V027 cluster were analysed using the HyPhy package for sequence evolution inference (Kosakovsky Pond et al. 2020). Three methods, Fixed Effects Likelihood (FEL), Single-Likelihood Ancestor Counting (SLAC) and Fast Unconstrained Bayesian AppRoximation (FUBAR) were applied to test for pervasive selection across the alignment. The sites under episodic diversifying selection (i.e. acting on a branch of a phylogenetic tree) were inferred by Mixed Effects Model of Evolution (MEME).

### Sample preparation for proteomic analysis

The specimens of salivary glands and glands of Leiblein for proteomic analysis were transported frozen to the laboratory and stored at −80°C. Each sample was used for both protein and intact peptide extraction, and three replicates of each tissue / species were analyzed resulting in a total of 12 analyzed samples. First, 500 μl of lysis buffer containing 2% sodium deoxycholate (SDC) in 100mM Tris (pH 8.5) preheated at 95°C was added to each specimen. Then the specimens were fragmented with scissors, and incubated at 95°C for 10 minutes. After the samples have cooled down they were subjected to sonication by Qsonica Q55 ultrasonic homogenizer (Qsonica, Newtown, CT, USA) at 80% amplitude using five series of five one-second-duration impulses. After the homogenization, the samples were centrifuges at 16000 × g for 10 min and the supernatants were transported to clean tubes. Cysteine reduction and alkylation were performed simultaneously by adding tris(2-carboxyethyl)phosphine (TCEP) up to 10 mM and chloroacetamide (CAM) up to 20 mM to the samples, following incubation at 56°C for 40 min. Meanwhile, the 10 kDa MWCO regenerated cellulose Amicon filters (Merck, Germany) were preconditioned by passing first 500 μl of 100mM Tris buffer and then the same Tris buffer containing 2% SDC trough each filter. The samples were applied to the filters and spun at 14000 × g until completely filtered. Then 200 μl of 0.5M NaCl was loaded onto each filter and spun at the same speed. Both portions of the flow-through from each sample were combined and stored for future peptide cleanup. The remaining filters containing the proteins in the upper chamber were washed twice with 500 μl of 100mM Tris buffer. Finally 200 μl of the same buffer were added to the upper chamber of each filter. The protein solutions were extensively mixed and the upper chambers were twisted upside down into new collecting tubes followed with the centrifugation of the filters to thoroughly collect the proteins from the filter membranes.

The resulting solutions of protein contained 100mM Tris buffer with the remnants of SDC allowing for protein measurement using BCA assay. Thirty micrograms of total protein were diluted up to 30 μl with the same Tris buffer. Then trypsin was added with the proportion of 1:50 and incubated overnight at 37°C. The reaction was terminated by the addition of trifluoroacetic acid (TFA) up to 1.5% to each sample resulting in consequential precipitation of SDC.

### Peptide cleanup

Trifluoroacetic acid (TFA) was added to all peptide samples obtained both after the filtration and trypsin digestion, up to 1.5% in order to remove SDC. Then three cycles of washing were performed. In each cycle, two volumes of ethyl acetate were added to the samples in order to dissolve the residual SDC precipitate and other unwanted contaminants. The samples were vortexed followed by quick centrifugation for 2 min at 6000 rpm (maximum speed in centrifuge BioSan Multi-spin MSV-6000, BioSan, Riga, Latvia), and the upper phase was discarded.

For the peptide desalting and cleanup the in-house made stage tips containing SDB-RPS membrane (Empore-3M, CDS Analytical, Oxford, PA, USA) were used. The tips were prepared according to (Rappsilber et al. 2003) with the use of 3 pieces of membrane in each tip. The samples were loaded into the tips and the tips were centrifuged at 1200 rpm (about 70 × g in the same centrifuge) until the solution has passed through the membrane. At the next step, 100 μl of 1% TFA covered by 50 μl of ethyl acetate were passed through the tips at the same speed in order to remove the remnants of SDC, and then washing was performed at the same speed with 100 μl 0.2% TFA. The peptides were eluted by passing 60 μl of 70% acetonitrile (CAN) with 5% NH_4_OH through the tips at the speed of 1000 rpm (about 50 × g). The peptide samples were then dried in the vacuum concentrator (Labconco, Kansas City, MO, USA).

### Liquid chromatography and tandem mass spectrometry (LC-MS/MS)

For the LC-MS analysis the samples were reconstituted in 0.1% TFA and loaded to a Acclaim PepMap 100 C18 (100 □m x 2 cm) trap column in the loading mobile phase (2% acetonitrile (ACN), 98% H_2_O, 0.1% TFA) at 10 μl/min flow and separated at 40°C on a 500 mm 75 µm inner diameter Thermo Scientific− Acclaim™ PepMap™ 100 C18 LC column with particle size 2 µm. Reverse-phase chromatography was performed with an Ultimate 3000 Nano LC System (Thermo Fisher Scientific), which was coupled to the Orbitrap Q Exactive HF mass spectrometer (Thermo Fisher Scientific) via a nanoelectrospray source (Thermo Fisher Scientific). the following chromatography conditions were used for the samples that underwent trypsin digestion: Water containing 0.1% (*v*/*v*) formic acid (FA) was used as mobile phase A and ACN containing 0.1% FA (*v*/*v*), 20% (*v*/*v*) H_2_O as mobile phase B. Peptides were eluted from the trap column with a linear gradient: 3–35% solution B (0.1% (v/v) formic acid, 80% (v/v) acetonitrile) for 105 min; 35-55% B for 18 min, 55-99% B for 0.1 min, 99% B during 10 min, 99-2%B for 0.1 min at a flow rate of 300 nl/min. After each gradient, the column was re-equilibrated with A for 10 min. Similar conditions were used for the samples containing intact peptides, but the total gradient time was 60 min. MS data was collected in DDA mode (TopN=15), with the following MS1 parameters: resolution120K, scan range350-1400, max injection time – 50 msec, AGC target – 3×10^6^. Ions were isolated with 1.2 m/z window, preferred peptide match and isotope exclusion. Dynamic exclusion was set to 30 s. MS2 fragmentation was carried out at 15K resolution with HCD collision energy set to 28, max injection time – 80 msec, AGC target – 1×10^5^. Other settings: charge exclusion-unassigned, 1, 6-8, >8.

### Bioinformatic integration of the mass spec data

The non-redundant catalogs of CDSs predicted from transcriptomic data of conspecific *Vexillum* specimens were used as databases for the proteomic search. Two different databases were built for each species. First database contained only mature toxin regions (Supp.Data3) of the transcripts of interest (i.e. putative venom components) and was used for the analysis of peptidomes in the samples of intact peptides. Second database contained all the predicted CDSs of a species, where the transcripts of interest contained a recognizable pattern in their sequence identifiers; this database was used for search of the spectra obtained after the trypsin digestions of the proteins.

All the .raw files were converted to .mzML format with ThermoRawFileParser (Hulstaert et al. 2020). The search engine IdentiPy v.0.3.3.16 (Levitsky et al. 2018) was used for proteins searches of data obtained after the trypsin digestion, followed by the post-search treatment and result filtration by Scavager v.0.2.4 (Ivanov et al. 2019). For the search, trypsin was chosen as a parameter and the number of allowed missed cleavages was set to 1. Mass accuracy for the precursor and the fragment ions were set to 10 ppm and 0.01 Da respectfully. Carbamidomethylation of Cys was set as a fixed modification, oxidation of Met, and deamidation of Gln and Asn - as variable modifications. The clusters of interest were filtered group-specifically with Scavager according to the target-decoy strategy with 1% false-discovery rate cut-off.

The .mzML files obtained for the samples with intact peptide extraction, were subjected to *de novo* peptide sequencing with PEAKS CMD (v. 1.0). The precursor and fragment mass accuracies were set to 10 ppm and 0.01 Da respectively, and no protease was selected, as there was no digestion performed for these samples. The results were filtered to at least 80% average confidence in peptide. The resulting peptides were mapped to the mature toxin sequences database, accounting for the identical masses of Leu and Ile.

