## Supplementary material for "Vexitoxins: a novel class of conotoxin-like venom peptides from predatory gastropods of the genus *Vexillum*": Kuznetsova et al. supplementary figures 1-9

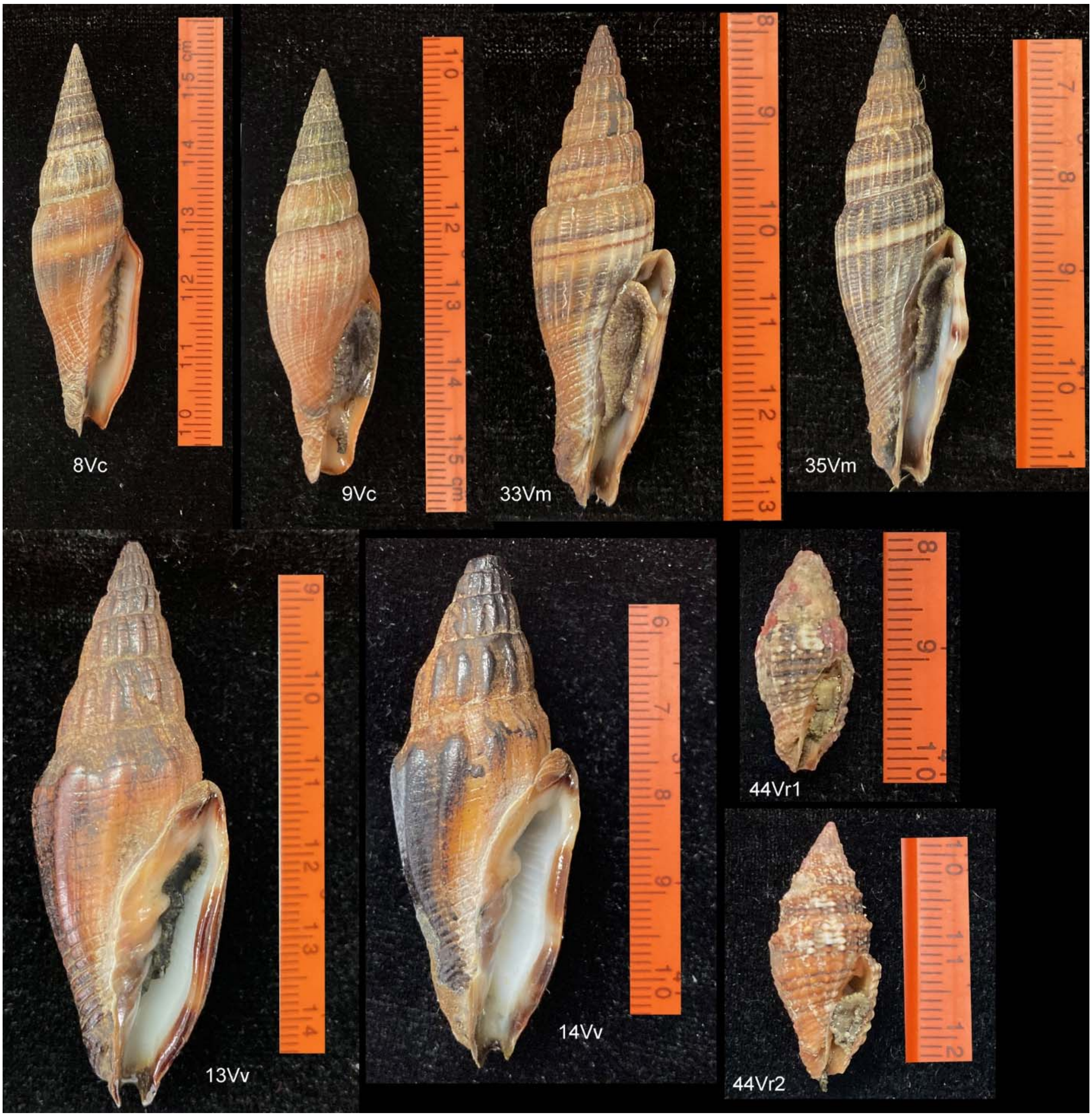


Figure S1. Figure S1. Vouchers of the *Vexillum* specimens dissected for RNA-Seq.

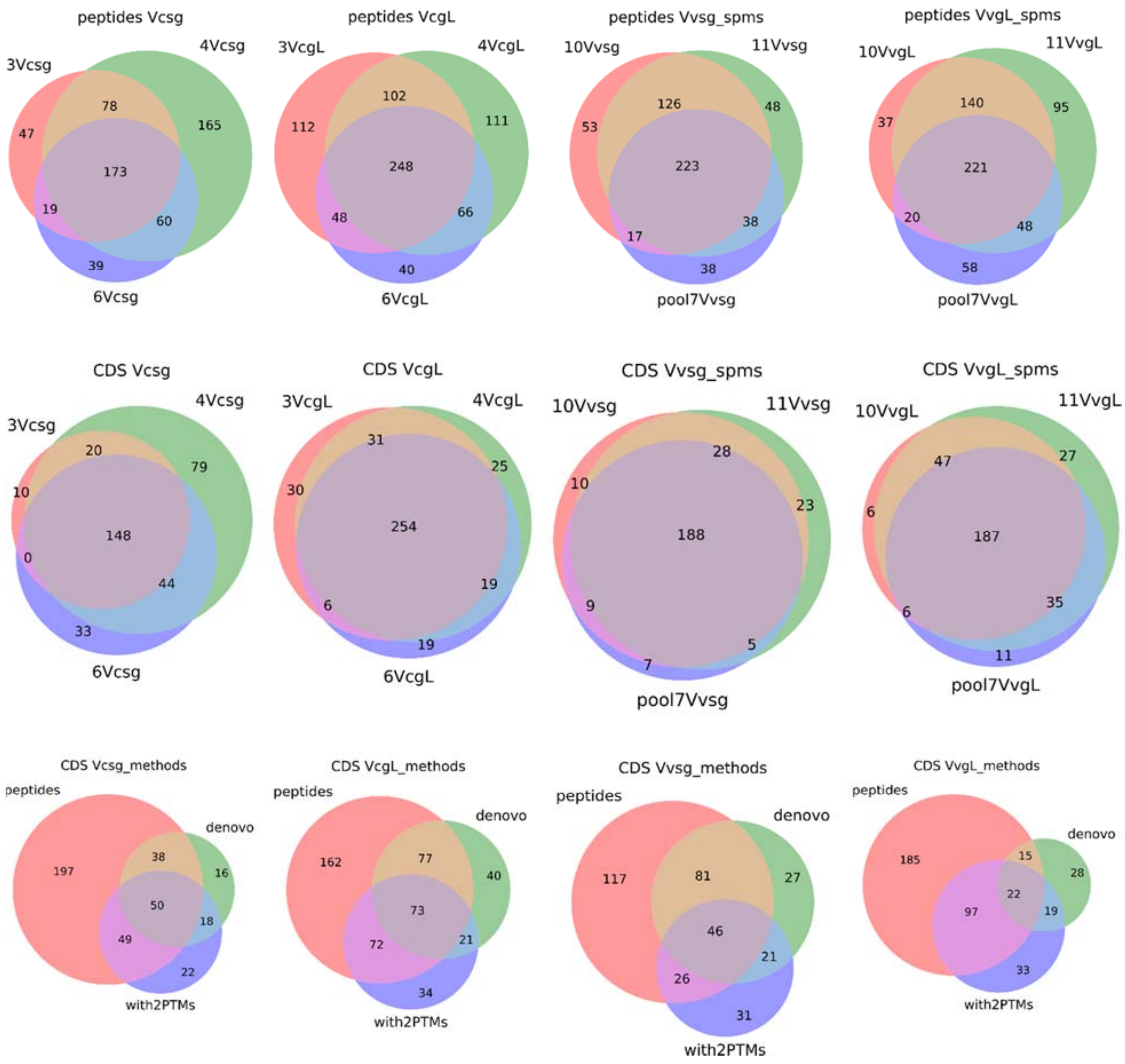


Figure S2. Venn diagrams showing contribution of profiled individualsand of different protocols of proteomic analysis to the detected number of peptides, and to the supported CDS (generated by transcriptomic analysis). **Top row**: overlap between the profiled replicates in the detected monoisotopic masses from the tripsin-lysed protein samples that match the search databases. **Second row**: overlap between the profiled replicates in the supported CDSs. **Bottom row**: contribution of different proteomic data sources to the support of query CDSs. Red: conventional peptide matching; Blue: peptide matching with mass mismatch corresponding to two PTMs (hydroxy-proline and carboxy-glutamic acid) allowed; Green *de novo* peptide sequencing based on the analysis of intact peptide fraction.

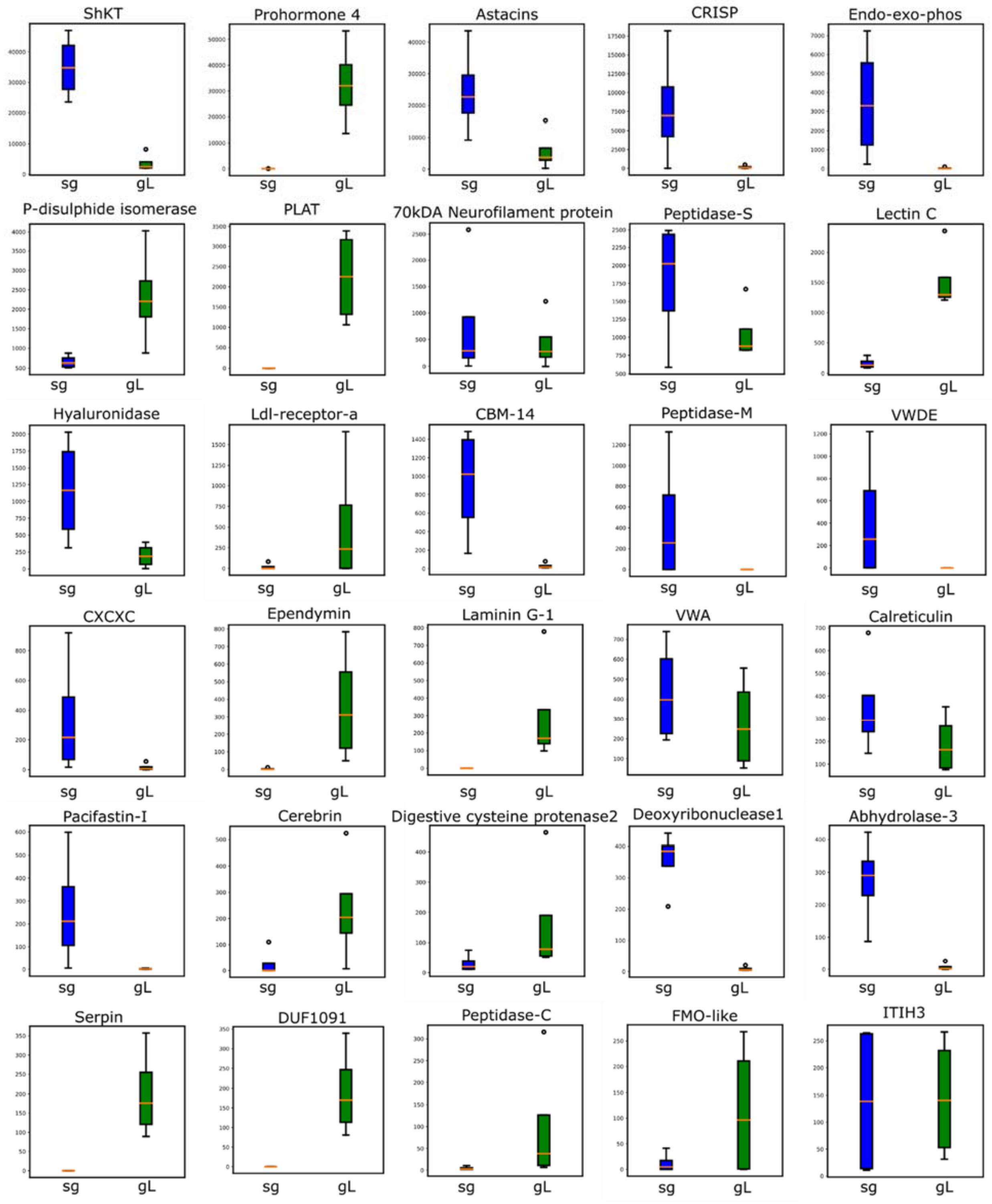


Figure S3. Boxplots of per-specimen summed TPM expression levels of 30 most highly expressed annotated transcripts clusters in sg (blue) and gL (green).

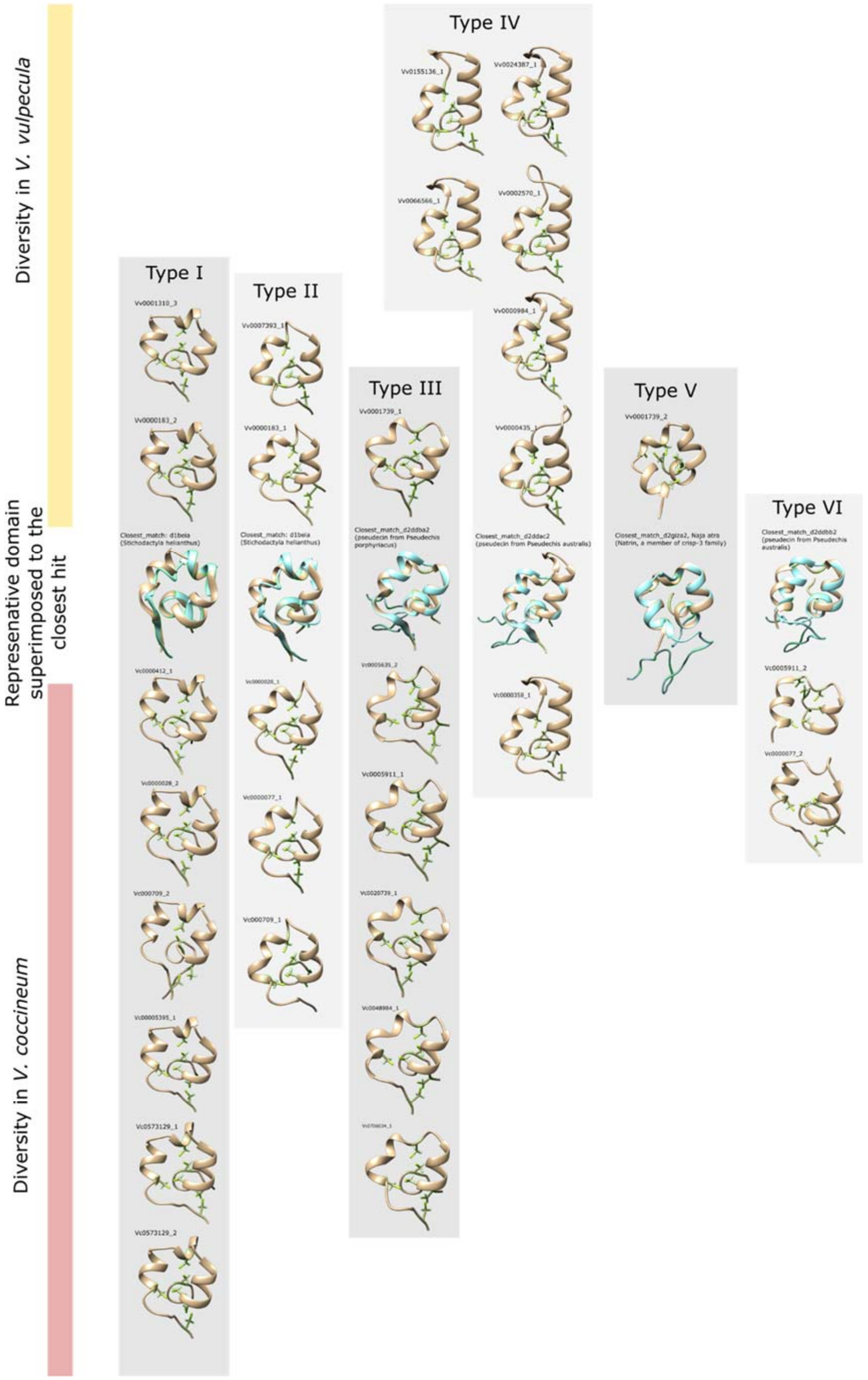


Figure S4. 3D structure of the 14 Vc and 11 Vv ShKT domains supported by the proteomic data. Representative members of each putative structural class superimposed by the closest structural hit detected by the RUPEE with SCOPe reference database. Title of each depicted domain like CDS-underscore-number of the domain from N-terminal.

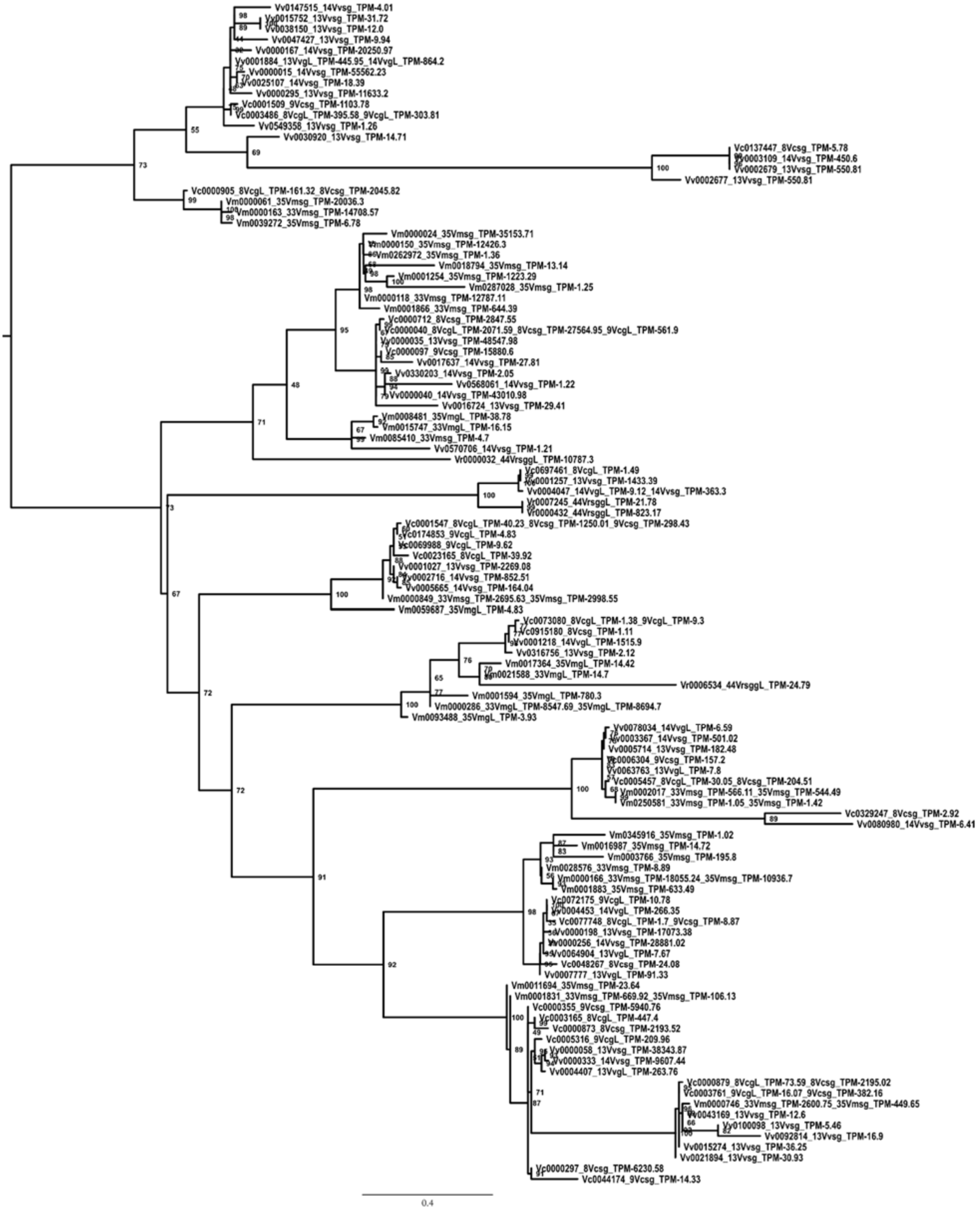

Figure S5. Mid-point rooted IQ tree-phylogenetic tree of the V001 cluster CDSs. Nodal support as ultrafast bootstrap.

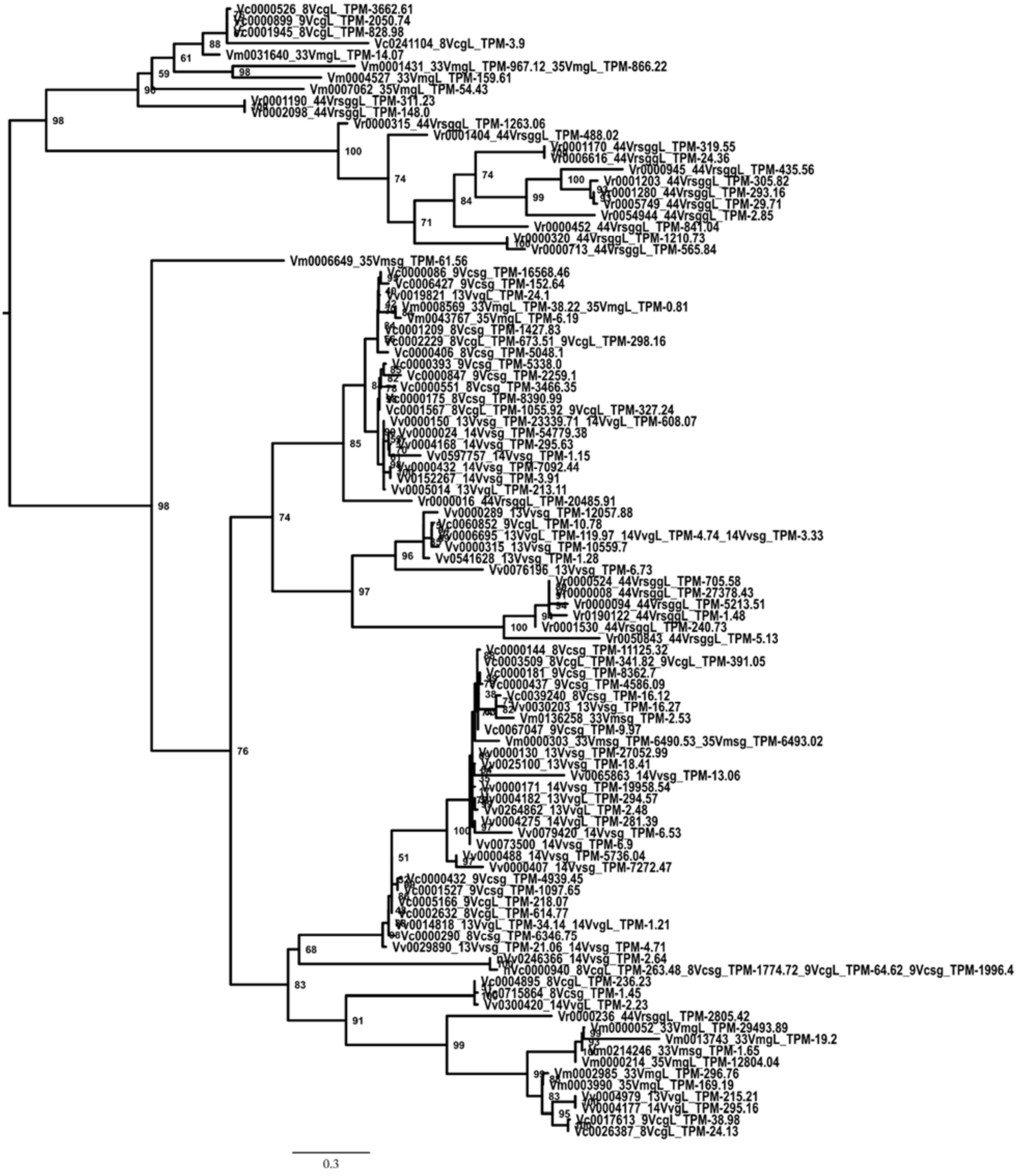


Figure S6. Mid-point rooted IQ tree-phylogenetic tree of the V002 cluster CDSs. Nodal support as ultrafast bootstrap.

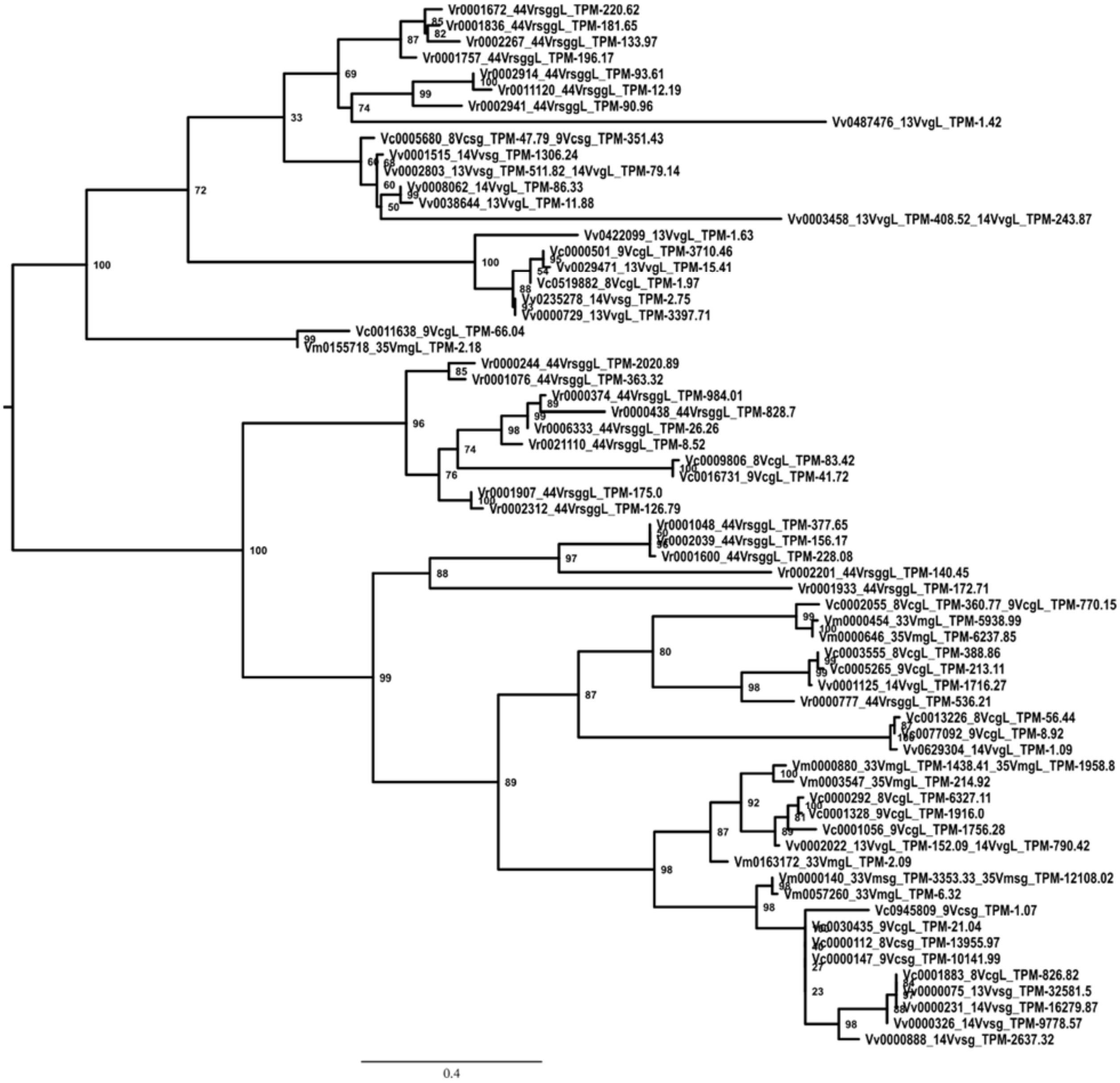


Figure S7. Mid-point rooted IQ tree-phylogenetic tree of the V004 cluster CDSs. Nodal support as ultrafast bootstrap.

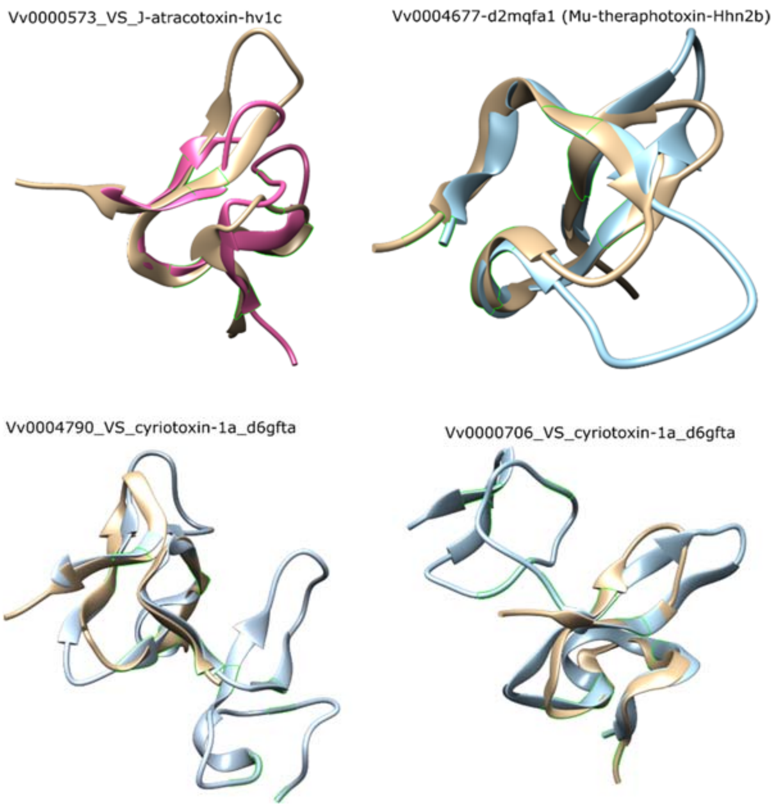


Figure S8. 3D structure of some putative toxins of *Vexillum*, containing a ICK motif (in pink or blue) in superposition to the most closely matching geometrically entry from a SCOPe database (beige).

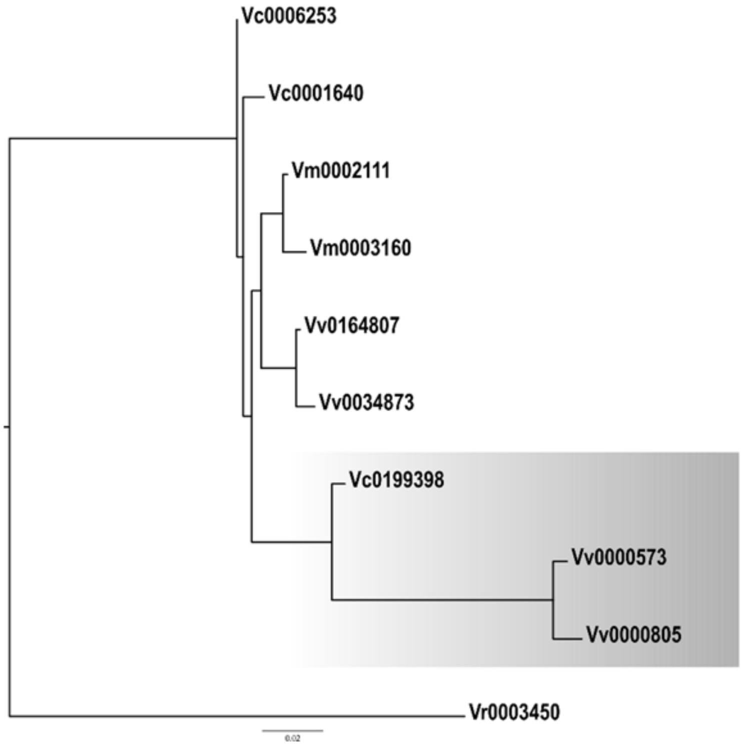


Figure S9. IQtree-based phylogenetic tree of the *Vexillum* V027 transcripts reconstructed from a codon alignment and rooted with the sequence Vr0003450 of *Vexillum crocatum*. The shorter orthogroup showed by grey shading.
